# Transcriptomic insights into developmental toxicity of per- and poly-fluoroalkyl substances (PFAS) in *Caenorhabditis elegans*: The potential key role of xenobiotic detoxification pathway

**DOI:** 10.1101/2025.11.21.689658

**Authors:** Zhenxiao Cao, Chenxi Zhou, Qing Zhao, Hua Du

## Abstract

Per– and polyfluoroalkyl substances (PFAS) are persistent environmental contaminants known to induce developmental toxicity across multiple species, yet the molecular mechanisms are still not fully understood. This study aims to evaluate the developmental toxicity of four long-chain legacy PFAS (PFOA, PFOS, PFNA, PFDA) and one short-chain alternative (PFBA) at environmentally relevant concentrations (1–5 μM) using the model organism *Caenorhabditis elegans*, with a focus on elucidating the underlying molecular mechanisms. Phenotypic analysis indicated that PFDA and PFOS significantly delayed development of worms, and reduced the number of fertilized eggs in the uterus. RNA-seq and subsequent bioinformatic analysis revealed strong impacts of PFDA and PFOS on physiological age. A core set of xenobiotic detoxification genes (e.g., *cyp-13A4*, *cyp-13A6*, and *cyp-13A7*), which were found to be primarily regulated by nuclear hormone receptors (NHR-102, NHR-85, NHR-28), showed consistent up-regulation upon PFAS exposure. Gene co-expression network analysis (WGCNA) further linked this detoxification gene signature to developmental impairment. Cross-species comparison using public databases identified several evolutionarily conserved detoxification genes that are associated with PFAS-induced developmental toxicity, among which CYP3A4 and its orthologs appear to be emerging biomarkers of PFAS exposure. Our findings demonstrate that activation of conserved xenobiotic detoxification pathways is a central transcriptomic signature of PFAS exposure, providing mechanistic insights into the structure-dependent developmental toxicity of this kind of pervasive pollutant.

Graphical Abstract

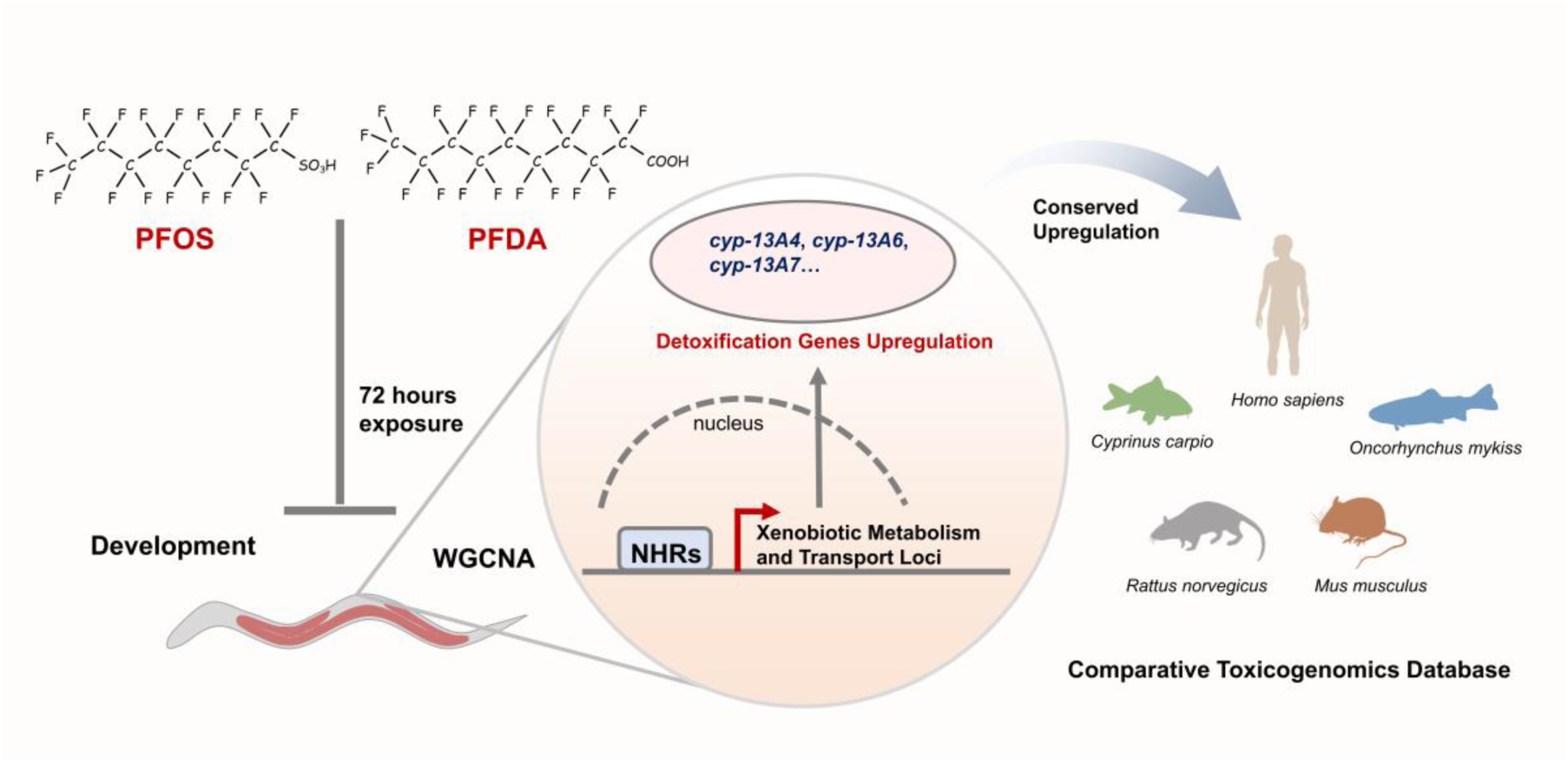

## 1. Introduction

Per– and polyfluoroalkyl substances (PFAS) are a class of synthetic chemicals having strong carbon-fluorine bonds. The remarkable thermal and chemical stability of PFAS has prompted their widespread use in commercial products since the late 1940s, including electronics, hydraulic fluids, and firefighting foams (Buck et al., 2011; Panieri et al., 2022; Ryu et al., 2021). Along with their production and consumption, PFAS have been progressively detected in the environment, even in remote regions such as the polar areas (He et al., 2025; Kelly et al., 2009). Environmental surveys have shown that PFAS concentrations in natural waters are typically in the ng/L-μg/L range, where they can reach up to mg/L levels near industrial sites (Kurwadkar et al., 2022; Liu et al., 2016). Owing to their high bioaccumulation potential, PFAS concentrations detected in human samples often exceed environmental levels (Rosato et al., 2024). Some of the most prevalent PFAS, such as perfluorooctane sulfonate (PFOS), have been reported as high as 10,400 ng/mL in human blood specimens (Zhou et al., 2014).

Previous studies have documented various toxic effects of PFAS, including metabolic dysregulation, hepatotoxicity, developmental disruption, and carcinogenic potential (Fenton et al., 2021). Developmental toxicity is of particular concern because PFAS exposure during critical windows—such as prenatal, early postnatal, or embryonic stages—may lead to long-lasting or irreversible biological outcomes (2021; Zhao et al., 2024). A landmark global meta-analysis has estimated that approximately 462,000 annual cases of low birth weight worldwide can be directly attributed to perfluorooctanoic acid (PFOA) exposure (Fan et al., 2022). Experimental studies have systematically demonstrated that PFAS exposure correlates with multiple developmental perturbations, including reduced body weight, delayed organogenesis, and malformations (ATSDR, 2021; Peritore et al., 2023). Accumulating evidence has suggested structure-toxicity relationships of PFAS, as shown by their toxic potential varying with structural attributes such as carbon chain length (Britton et al., 2024; Feng et al., 2023).

During the past two decades, the developmental toxicity of two long-chain PFAS (PFOA and PFOS) has been thoroughly examined. As long-chain PFAS are progressively restricted or banned globally under regulatory frameworks such as the Stockholm Convention, the manufacturing, utilization, and environmental release of their short-chain alternatives have increased substantially (UNEP; Xiao et al., 2022). Notably, perfluorobutanoic acid (PFBA), a PFOA substitute, has shown remarkable environmental prevalence in certain contaminated regions (Cai et al., 2020; Paige et al., 2024). Although short-chain PFAS are generally perceived as less toxic than their long-chain counterparts, their toxicological profiles and underlying mechanisms remain poorly investigated compared with legacy PFAS (Wang et al., 2024). This may hinder effective regulation of emerging PFAS contaminants, leading to potential public health and ecological risks.

In our earlier work, the multi-generational effects and developmental toxicity of four legacy PFAS have been examined. The findings show a minor contribution of genetic damage to the reproductive and developmental toxicity of PFAS (Cao et al., 2024). Alterations in gene expression are a critical biological response to environmental stress (Johnson et al., 2022). Prior transcriptomic studies have documented disturbance effects of PFAS on core developmental and metabolic signaling pathways, suggesting possible connections between transcriptomic dysregulation and PFAS-induced developmental toxicity (Yao et al., 2019; Zhou et al., 2025). In this study, we set out to investigate developmental toxicity of four long-chain PFAS (PFOA, PFOS, perfluorononanoic acid [PFNA], and perfluorodecanoic acid [PFDA]) and one short-chain PFAS (PFBA) in *C. elegans*, and to explore commonly dysregulated genes in response to PFAS exposure. Among all five tested PFAS, environmentally relevant concentrations (1–5 μM) of PFDA and PFOS exhibited significantly greater impacts on the development of worms. Using RNA-seq and comprehensive bioinformatic analysis, PFAS-induced transcriptomic alterations were examined, and differentially expressed key genes were identified. The results revealed a robust transcriptome signature associated with PFAS-induced developmental toxicity, as shown by pronounced upregulation of xenobiotic detoxification genes. Cross-species comparative analysis further demonstrated an evolutionarily conserved subset of these genes, while cytochrome P450 (CYP) gene CYP3A4 orthologs were found to be potential biomarkers of PFAS exposure. In sum, our findings advance mechanistic understanding of developmental toxicity induced by both legacy PFAS and the emerging alternatives.

## 2. Materials and Methods

### 2.1. Chemicals and C. elegans growth conditions

PFOA (CAS No. 335-67-1), PFNA (CAS No. 375-95-1), PFDA (CAS No. 335-76-2), and PFBA (CAS No. 375-22-4) were obtained from Sigma-Aldrich (St. Louis, MO, USA). PFOS (CAS No. 1763-23-1) was purchased from Dr. Ehrenstorfer (Augsburg, Germany). Stock solutions of these chemicals (50–100 mM) were prepared in dimethyl sulfoxide (DMSO; Sangon Biotech, Shanghai, China) and stored at 4 °C until use.

The wild-type *C. elegans* N2 Bristol strain was obtained from Caenorhabditis Genetics Center (CGC; Minneapolis, MN, USA). Worms were maintained in the dark at 20 °C on Nematode Growth Media (NGM) plates with *Escherichia coli* OP50 seeded as the food source. Synchronized L1 larvae were obtained by treating gravid adult hermaphrodites with a bleaching solution (5% NaClO and 5 M NaOH at a 2:1 volume ratio).

### 2.2. Exposure experimental design

The PFAS stock solutions were diluted to treatment concentrations (1–5 μM) in M9 buffer. Synchronized L1 larvae were exposed to PFAS in 24-well plates. For the daily brood size assay, the exposure duration was 60 hours, while a 72-hour exposure period was used for all other experiments. The exposure medium was supplemented with an appropriate amount of *Escherichia coli* OP50 as a food source.

### 2.3. Oocyte and fertilized egg enumeration assay

Following exposure, worms were washed three times and stained with 25 μg/mL acridine orange (AO; Sangon Biotech, Shanghai, China) for 1 h. After staining, worms were transferred to fresh NGM plates and allowed to recover for 45 minutes to expel intestinal AO, thereby preventing interference with oocyte quantification. Subsequently, worms were anesthetized using levamisole (J&K Scientific, Beijing, China), and the numbers of both fertilized eggs in the uterus and oocytes in the gonads were counted under an inverted fluorescence microscope (DMi8, Leica, Wetzlar, Germany). At least 15 worms per experimental group were analyzed for fertilized eggs. Oocytes in the gonad arm proximal to the tail were counted in at least 10 worms per experimental group.

### 2.4. Developmental stage analysis

Worms were immobilized with levamisole and examined under the bright-field mode of an inverted microscope to determine developmental stages and quantify the proportion of worms at each developmental stage. The staging criteria were based on vulval morphology, as described previously (Porta-de-la-Riva et al., 2012).

Approximately 100 worms were analyzed per group.

### 2.5. Daily brood size assay

Brood size was quantified daily over a five-day period following the cessation of exposure, as previously described (Cao et al., 2024). Detailed procedures are provided in Text S1.

### 2.6. RNA extraction and sequencing

Following exposure, worms were washed three times with M9 buffer to remove residual *E. coli* OP50. After adding two zirconia beads to each sample, the mixtures were frozen at –80 °C for 10 minutes and mechanically homogenized using a pre-chilled bead mill for 3 minutes. Total RNA was extracted using TRIzol reagent (Invitrogen, Waltham, Massachusetts, USA) following the manufacturer’s protocol. Extracted RNA samples were sent to Biozeron Biotech Ltd. (Shanghai, China) for quality control and sequencing. High-quality RNA samples (OD260/280 = 1.8–2.2, OD260/230 ≥ 2.0, RIN ≥ 6.5, 28S:18S ≥ 1.0, total RNA > 2 μg) were selected for library preparation. Strand-specific RNA-seq libraries were constructed using the KAPA Stranded mRNA-Seq Library Preparation Kit, followed by paired-end sequencing (150 bp × 2) on an Illumina NovaSeq 6000 platform (Illumina, San Diego, CA, USA).

### 2.7. RNA-sequencing data analysis

Raw FASTQ files were processed using Trimmomatic (v0.39) for quality control. Cleaned reads were aligned to the *C. elegans* reference genome WS284 using HISAT2 (v2.2.1). Gene expression quantification was performed with FeatureCounts (v2.0.6) to generate the raw count matrix. Differential expression analysis was conducted using the DESeq2 (v1.46.0) package in R (v4.4.2) (Love et al., 2014), with significantly differentially expressed genes (DEGs) defined as those with |log₂FoldChange| > 1 and adjusted p value < 0.05.

Nematode age estimation and correction for age effects in differential expression analysis were performed using the RPAToR (v1.2.0) R package (Bulteau and Francesconi, 2022). We staged the samples with the “Cel_larv_YA” of the wormRef package using 1,000 interpolated time points. Principal component analysis (PCA) was performed using the OmicShare platform (https://www.omicshare.com/). Gene set enrichment analysis of DEGs was performed via the WormCat web platform using the “whole genome v2” annotation type (Higgins et al., 2022). KEGG pathway enrichment analysis was conducted with the clusterProfiler R package (v4.14.3) (Yu, 2024). Protein–protein interaction (PPI) network of detoxification genes was constructed in Cytoscape (v3.10.3) using the stringApp plugin (Su et al., 2014). Hub genes were identified via the cytoHubba plugin with the Maximal Clique Centrality (MCC) algorithm. Transcription factor activity was evaluated using the CelEST Shiny app (v1.0) on RNA-seq raw counts with developmental age correction (Perez, 2025). WGCNA was performed using the WGCNA R package (v1.73) on TPM-normalized data, with a soft threshold of 12 to construct a signed network. Further details on the WGCNA procedure are provided in Text S1.

For cross-species analysis, *C. elegans* genes were mapped to human orthologs using the WormBase database. For genes without annotated orthologs in WormBase, the DIOPT R package (v0.2.1) was used to query the DIOPT online tool (v9.0) in “best match” mode (https://www.flyrnai.org/diopt). Gene expression trends across species were compared using data from the Comparative Toxicogenomics Database (CTD) and the results were further filtered in R (v4.4.2) to identify consistently regulated genes (Davis et al., 2023). Data visualization was conducted with the ggplot2 (v3.5.1) R package.

### 2.8. RNAi treatment

The RNA interference (RNAi) experiment was carried out with modifications based on previously established protocols (Hammell and Hannon, 2012). A detailed description is provided in Text S1.

### 2.9. Real-time quantitative PCR

Total RNA was reverse transcribed into cDNA using the cDNA Synthesis SuperMix (TransGen Biotech, Beijing, China). For qPCR amplification, reactions were performed using PerfectStart Green qPCR SuperMix (TransGen Biotech, Beijing, China) on a LightCycler 96 real-time PCR system (Roche, Basel, Switzerland). All qPCR experiments were performed according to the manufacturer’s instructions. Relative gene expression levels were calculated using the 2^−ΔΔCt^ method and normalized to the *tba-1* gene in each sample. Primer sequences are listed in Table S1.

### 2.10. Statistical Analysis

Data were derived from at least three independent experiments. Homogeneity of variances was assessed using Levene’s test. Statistical comparisons among multiple groups were performed using one-way analysis of variance (ANOVA) followed by Bonferroni correction for multiple comparisons. For comparisons between two groups, a two-tailed Student’s t-test was performed. The significance of overlapping differentially expressed genes between experimental groups was evaluated using the hypergeometric test. All statistical analyses were conducted using R (v4.4.2).

## 3. Results

### 3.1. Impacts of PFAS on development of C. elegans

Nematode development undergoes five sequential stages: L1, L2, L3, L4, and adult stage. To investigate the dose-response relationship for developmental toxicity of five PFAS (Figure 1A), the proportions of *C. elegans* at each developmental stage were assessed after a 72-hour exposure to serial concentrations (1, 3, 5, 7, and 9 μM) of PFAS. The results demonstrated that PFBA, PFOA, and PFNA did not alter developmental stage of worms. In contrast, PFDA exposure markedly delayed worm development at concentrations ≥3 μM (Figure 1B). The proportion of immature worms (L1-L4) dose-dependently increased to 8.52% (3 μM), 25.97% (5 μM), 93.73% (7 μM), and 99.18% (9 μM) following PFDA treatment, compared to 3.11% in the control group. Similarly, PFOS at concentrations ≥5 μM significantly hindered worm development, as shown by 11.11% (5 μM), 33.91% (7 μM), and 89.38% (9 μM) immature worms, versus 2.20% in the control (Figure 1B). According to these results, PFAS ranging from 1 to 5 μM were subsequently applied to further examine their developmental toxicity.

**Figure 1.**
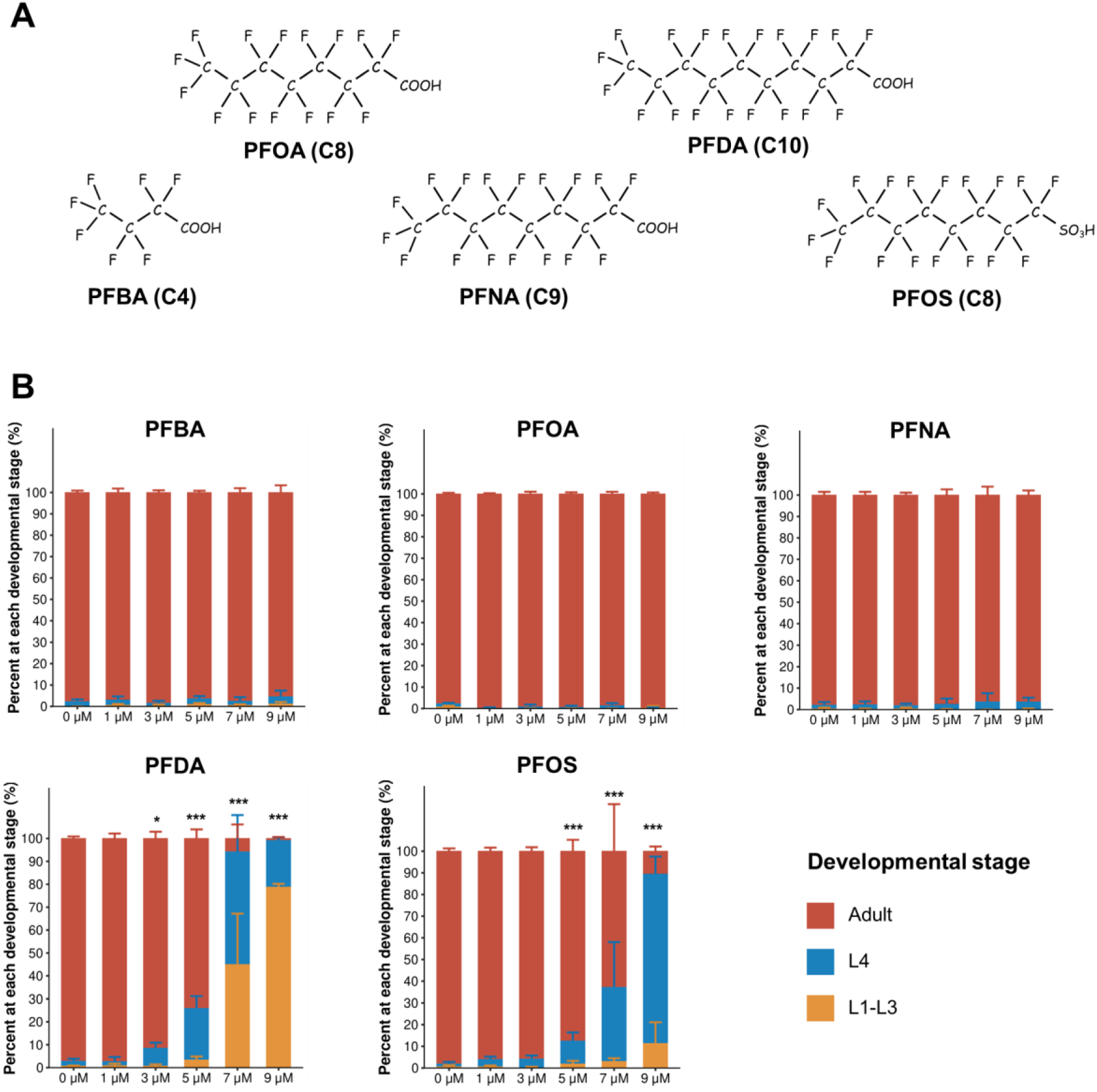
Five PFAS tested in this study and their effects on *C. elegans* development. (**A**) Chemical structures of PFBA, PFOA, PFNA, PFDA and PFOS. Numerals in parentheses denote carbon chain lengths. (**B**) Developmental toxicity of PFAS. Synchronized L1-stage nematodes were exposed to PFAS for 72 h, followed by quantification of proportions at each developmental stage. Error bars represent SEM. Differences between PFAS-treated groups and the control were analyzed by chi-square test with Bonferroni correction. \**p* < 0.05, \*\*\**p* < 0.001.

### 3.2. Effect of PFAS on reproductive fitness of C. elegans

After a 72-hour exposure, total brood size was not significantly affected by any of PFAS tested (Figure S1). However, 3 μM PFDA pronouncedly reduced offspring numbers per worm on both day 1 and day 2 (day 1: 3.37 vs. 26.27, *p* = 0.027; day 2: 80.40 vs. 111.86, *p* = 0.03; Figure 2A). The reproductive impairment was more pronounced under 5 μM PFDA treatment, as revealed by 0.85 offspring count on day 1, and 55.68 offspring count on day 2 (*p* = 0.03 and *p* = 3.7 × 10⁻^5^, respectively).

**Figure 2.**
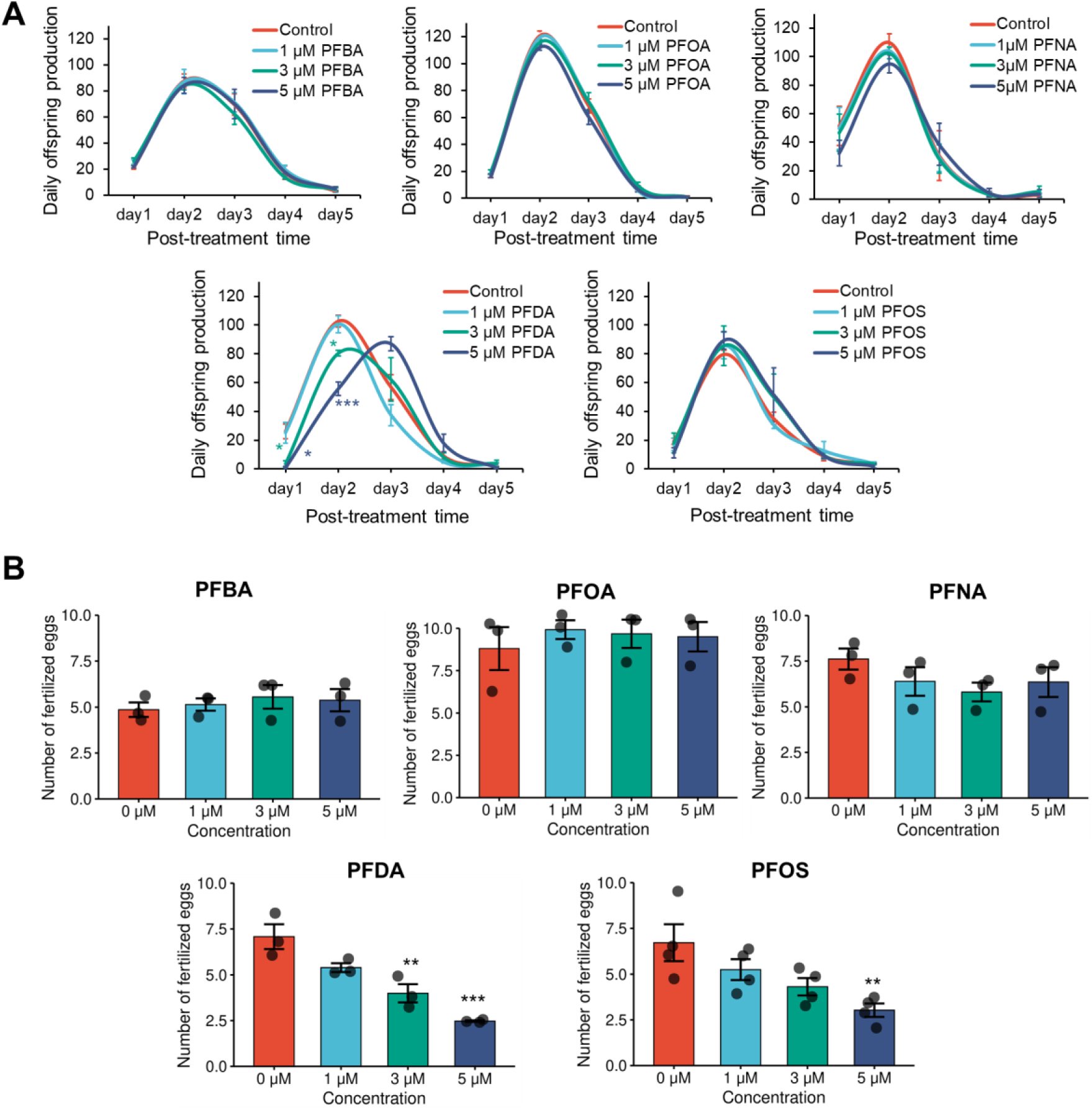
Effects of five PFAS on *C. elegans* reproduction. After exposure of synchronized L1 larvae to PFAS. (**A**) daily offspring production over 5 days and (**B**) the number of fertilized eggs in the uterus were quantified. Error bars represent SEM. Statistical differences between PFAS-treated groups and the control were analyzed using one-way ANOVA with Bonferroni correction: \**p* < 0.05, \*\**p* < 0.01, \*\*\**p* < 0.001.

Analysis of the number of fertilized eggs in the uterus also indicated reproductive toxic effects of PFDA and PFOS. For PFDA, 3 μM and 5 μM exposures led to 43.64% (*p* = 0.006) and 65.11% (*p* = 4.4 × 10⁻⁴) declines in the number of fertilized eggs, respectively (Figure 2B). Similarly, fertilized egg counts were reduced by 35.79% and 54.84% (*p* = 0.009) upon 3 μM and 5 μM PFOS treatment. However, none of the five tested PFAS altered oocyte counts (Figure S2).

### 3.3. PFAS exposure altered physiological age of worms

Transcriptome sequencing was performed following 5μM PFAS exposure. Principal component analysis (PCA) of the gene expression data (FPKM) revealed high reproducibility across the three biological replicates in each group (Figure 3A). Moreover, the PFDA– and PFOS-treated groups showed distinct transcriptomic profiles from those of the other PFAS-treated groups and the control group.

**Figure 3.**
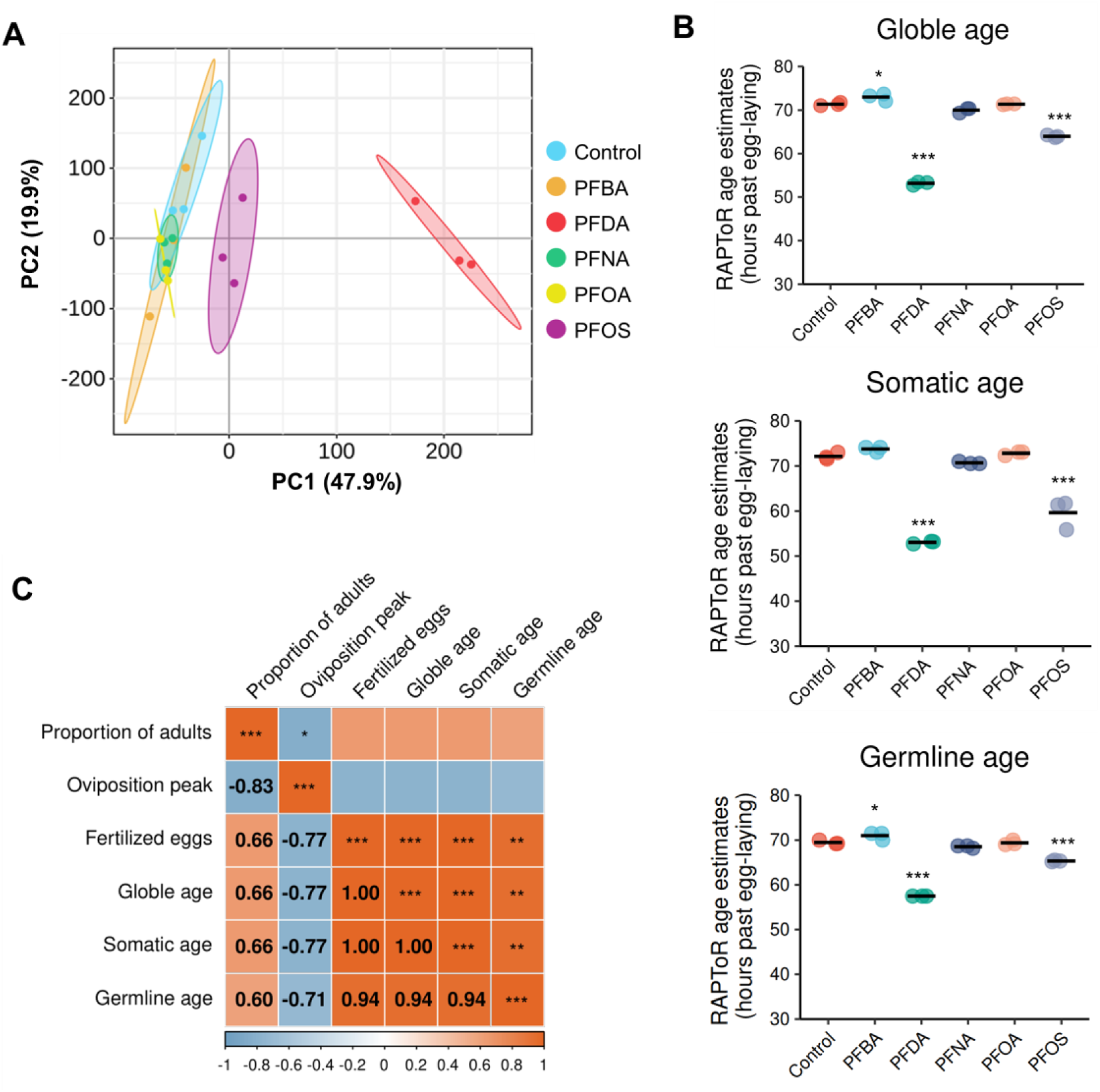
Transcriptomic profiling and physiological age estimation in *C. elegans* following PFAS exposure. (**A**) Principal component analysis of mRNA expression profiles (**B**) Global, somatic, and germline age estimates based on transcriptomic data using the RAPToR R package. Each point represents a biological replicate, with the mean line indicating the average of three replicates. Statistical significance was determined by one-way ANOVA followed by Bonferroni adjustment for multiple comparisons between control and treatment groups. (**C**) Spearman correlation analysis between RAPToR-estimated ages and reproductive/developmental toxicity endpoints. Color intensity maps to correlation coefficients. \**p* < 0.05, \*\**p* < 0.01, \*\*\**p* < 0.001.

RAPToR analysis revealed that PFDA– and PFOS-treated worms exhibited significantly younger physiological ages, estimated at 53.17 h (*p* = 1.47×10^-13^) and 63.99 h (*p* = 6.14×10^-9^) post egg-laying, compared with 71.38 h in control group (Figure 3B). In contrast, PFBA exposure resulted in a pronounced increase in physiological age (73.01 h, *p* = 0.03). Germline and somatic ages were then identified using tissue-specific gene sets (Figure 3B). While PFBA treatment did not alter somatic age, it led to a remarkable increase in germline age (71.01 h vs. 69.49 h in the control group, *p* = 0.03). The influences of the other four PFAS compounds on both germline and somatic ages were consistent with their impacts on global physiological age. It is worth noting that RAPToR-estimated physiological age showed a strong positive correlation with fertilized egg counts (Figure 3C).

### 3.4. Comparative analysis of DEGs across PFAS treatments

To further elucidate PFAS-induced changes in gene expression, DEGs were identified using a standard DESeq2 pipeline, with the thresholds |log2FC| > 1 and adjusted *p*-value < 0.05 relative to the control. PFDA and PFOS treatment groups yielded markedly higher numbers of DEGs (8904 and 2826, respectively) compared with the other three PFAS treatment groups (ranging from 5 to 154 DEGs), with the majority of DEGs being upregulated (Figure 4A-C).

**Figure 4.**
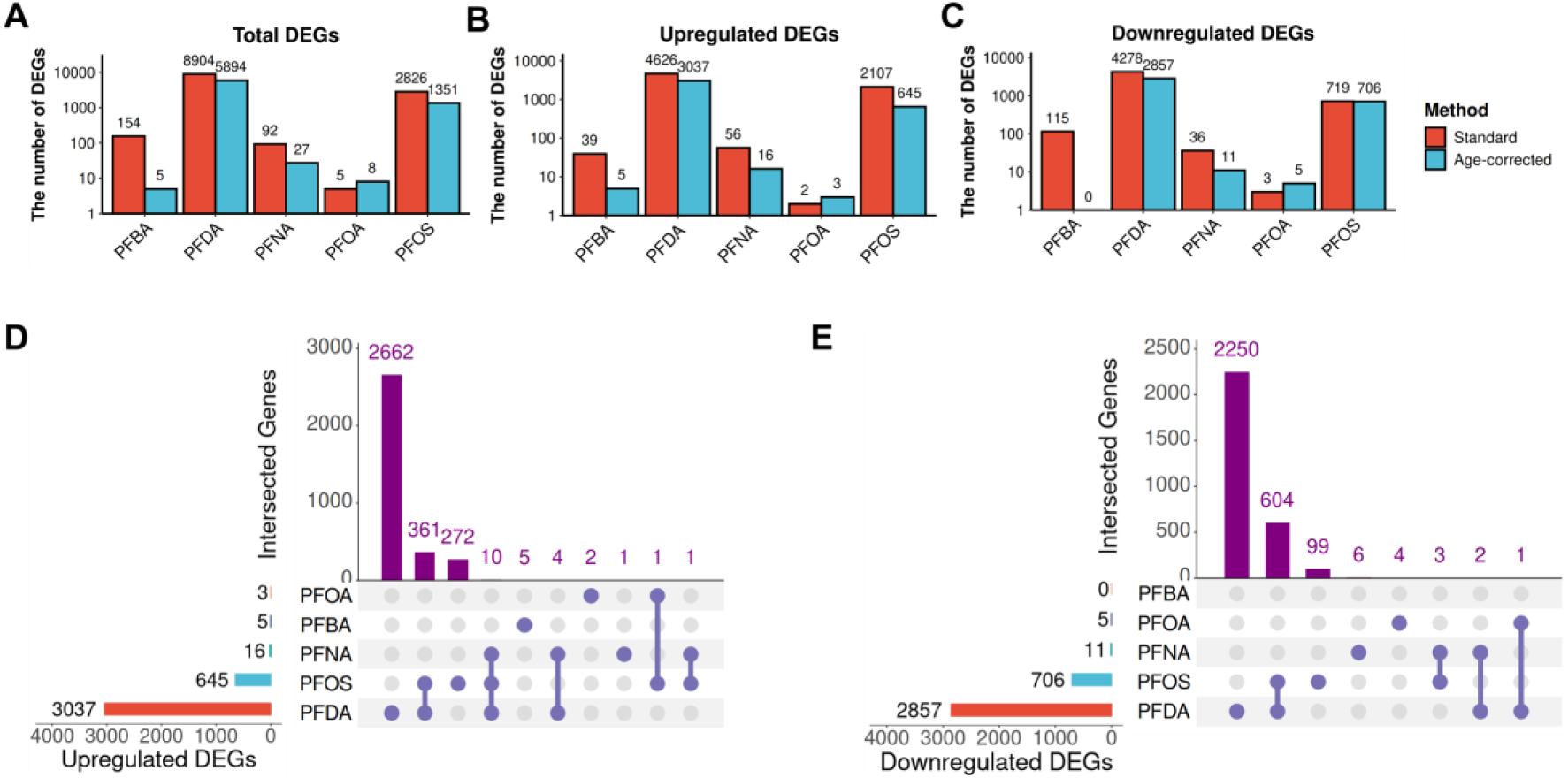
Number and overlap of DEGs across PFAS treatments. (**A–C**) The number of (**A**) total, (**B**) up-regulated, and (**C**) down-regulated DEGs. Standard DEGs were identified via the conventional DESeq2 pipeline, while age-corrected DEGs were generated using the RAPToR R package. (**D–E**) UpSet plots illustrating overlaps among (**D**) up-regulated and (**E**) down-regulated age-corrected DEGs across treatments.

By using the RAPToR R package to quantify age-driven expression changes, we found that gene expression alterations in PFOS– and PFDA-treated groups were highly correlated with the expected developmental expression changes calculated from matching time points in the reference (r = 0.725 and 0.783, respectively; Figure S3), indicating a strong contribution of developmental age to the observed transcriptomic changes. We therefore removed age as a confounding factor and identified PFAS-specific perturbations on gene expression using RAPToR. The number of age-corrected DEGs was generally reduced compared to that from the standard pipeline (Figure 4A-C). Hypergeometric tests on overlaps of age-corrected DEGs across treatment groups showed significant concordance among PFAS-induced DEGs, and the largest overlap was found between PFDA and PFOS groups (*p* < 0.001, Figure 4D-E; Figure S4).

Volcano plots of the age-corrected DEGs (Figure 5A-E) showed that cytochrome P450 (CYP) genes were among the top five up-regulated genes in PFDA-, PFNA-, and PFOS-treated groups. Specifically, PFDA upregulated *cyp-13A4* and *cyp-13A8*; PFNA upregulated *cyp-13A6*, *cyp-13A7* and *cyp-34A10*; and PFOS upregulated *cyp-13A4*, *cyp-13A6*, and *cyp-13A7*. A similar pattern was observed in DEGs identified using the standard pipeline (Figure S5A).

**Figure 5.**
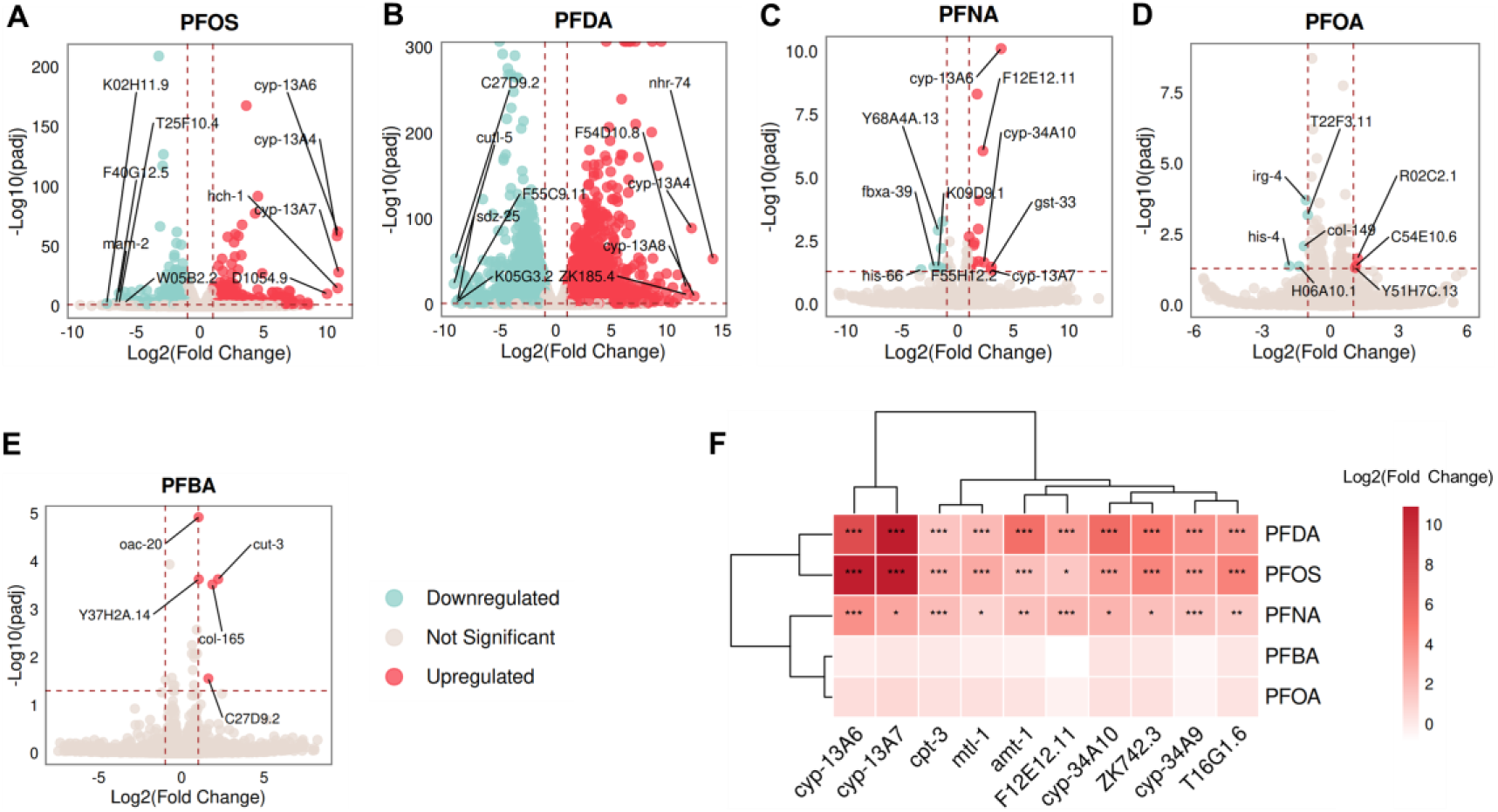
Distinct and shared age-corrected DEGs across PFAS treatments. (**A–E**) Volcano plots of age-corrected DEGs for each PFAS treatment. The top five most significantly upregulated and downregulated genes in each group are labeled. (**F**) Heatmap displaying age-corrected DEGs that were consistently and significantly up– or down-regulated in at least three treatment groups. * adjusted *p* < 0.05, ** adjusted *p* < 0.01, *** adjusted *p* < 0.001.

We further identified genes consistently upregulated or downregulated in at least three treatment groups to identify common PFAS-responsive genes. This analysis revealed 10 consistently upregulated genes, including four CYP genes: *cyp-13A6*, *cyp-13A7*, *cyp-34A9*, and *cyp-34A10* (Figure 5F). In contrast, the standard pipeline identified 68 such genes, including seven CYP genes (Figure S5B). The upregulation of these genes were further validated by RT-qPCR (Figure S6). By conducting RNA interference (RNAi) targeting *cyp-13A4* and *cyp-13A6*, we found that *cyp-13A6* RNAi partially restored fertilized egg counts in PFDA-treated worms from 26.45% to 33.84% of control levels (*p* = 0.0499, two-tailed t-test) (Figure S7), suggesting the potential modulatory effects of CYP genes on PFAS-induced developmental toxicity.

### 3.5. Enrichment analysis of PFAS-induced DEGs

Both age-corrected DEGs and standard DEGs were subjected to WormCat enrichment analysis (a web-based tool specialized for *C. elegans* gene enrichment), and KEGG pathway enrichment analysis (Figure 6A-B, Figure S8-S10). The WormCat analysis revealed that among the upregulated age-corrected DEGs, terms associated with xenobiotic detoxification, including solute carriers (SLCs), ATP-binding cassette (ABC) transporters, UDP-glucuronosyltransferases (UGTs), glutathione S-transferases (GSTs), short-chain dehydrogenases (SDRs), and CYPs, were significantly enriched in at least one treatment group (Figure 6A). Notably, CYP genes were consistently enriched across PFOS-, PFDA-, and PFNA-exposed groups (*p* < 0.05). KEGG pathway analysis further demonstrated that xenobiotic metabolism, detoxification, and transport pathways—including those associated with CYPs, GSTs, and ABC transporters—were significantly enriched in PFOS– and/or PFDA-treated groups among upregulated age-corrected DEGs (Figure 6B). Similar patterns of enrichment were observed in standard DEGs (Figure S9A, S10A). Among all age-corrected DEGs, there were 226, 66, and 5 xenobiotic detoxification genes identified in PFDA-, PFOS-, and PFNA-treated groups, respectively. In contrast, no xenobiotic detoxification genes were detected in the other two PFAS-treated groups (Figure S11).

**Figure 6.**
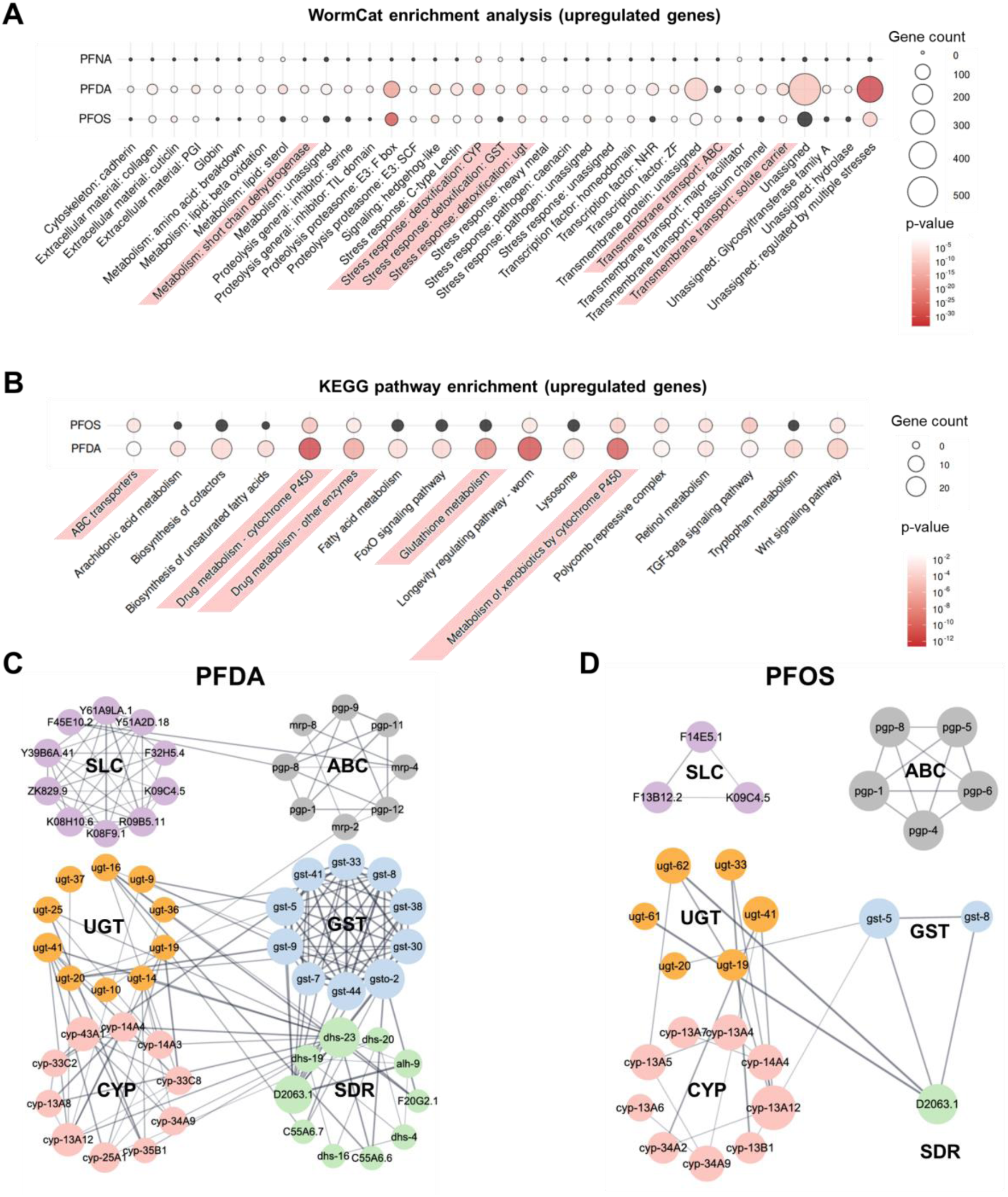
Gene enrichment analyses and PPI networks for age-corrected DEGs. (**A–B**) WormCat and KEGG pathway enrichment analyses for up-regulated DEGs. Only terms with *p* < 0.001 in at least one treatment group are displayed; black bubbles represent non-significant terms. (**C–D**) PPI networks of hub DEGs within WormCat categories associated with xenobiotic detoxification and metabolism under (**C**) PFDA and (**D**) PFOS treatment. Hub genes were identified from each category using the CytoHubba plugin in Cytoscape, with the top 10 genes by MCC score being plotted when available.

Using Cytoscape and its plugins, hub genes associated with xenobiotic detoxification within the age-corrected DEG sets of PFOS and PFDA were identified, and PPI networks were constructed (Figure 6C-D). The data revealed strong interactions among gene modules including UGTs, GSTs, CYPs, and SDRs, whereas interactions between SLC/ABC transporter families and these modules were relatively weak.

### 3.6. Gene co-expression modules associated with PFAS-induced developmental toxicity

We employed WGCNA to investigate associations between PFAS-induced transcriptomic alterations and corresponding phenotypic changes, as well as RAPToR-estimated physiological ages. A scale-free co-expression network was constructed with a soft threshold power of 12 (Figure S12). Gene expression profiles were clustered into 12 modules, each containing 44-3044 genes (Figure 7A-B). Spearman correlation analysis revealed that the greenyellow, purple, brown, blue, and pink modules were significantly correlated with all traits assessed (Figure 7A). Among them, only the greenyellow module exhibited a strong negative correlation with RAPToR-estimated ages (r = –0.95, – 0.96, and –0.97 for global, somatic, and germline ages, respectively; Figure 7A), which contains the largest number of xenobiotic detoxification genes (n = 111), including members of ABC transporter, SLC, UGT, GST, SDR, and CYP families (Figure 7B). We further analyzed module eigengene (ME) expression patterns across all modules (Figure 7C, Figure S13). In the greenyellow module, ME expression in PFDA-treated group showed the most significant elevation as compared with control group (*p* = 3.49×10^-12^), followed by that in PFOS-treated group (*p* = 3.95×10^-12^, one-way ANOVA with Bonferroni correction). Both KEGG pathway and WormCat enrichment analysis demonstrated pronounced enrichment of xenobiotic detoxification-related terms in the greenyellow module (Figure 7D-E). Intriguingly, PFOS– and PFDA-treated groups exhibited concurrent enrichment of specific WormCat terms, including “Transmembrane protein: unassigned”, “Cytoskeleton: microtubule: tau tubulin kinase”, “Signaling: phosphatase: Y”, “Signaling: Y kinase”, “Unassigned: membrane spanning domain”, and “Major sperm protein”, all of which were observed in both downregulated age-corrected DEGs and the greenyellow module (Figure 7D, Figure S8A). For KEGG pathway, “Cysteine and methionine metabolism” and “Cytoskeleton in muscle cells” terms were enriched in the downregulated age-corrected DEGs of both PFOS and PFDA groups, as well as in the greenyellow module (Figure 7E, Figure S8B).

**Figure 7.**
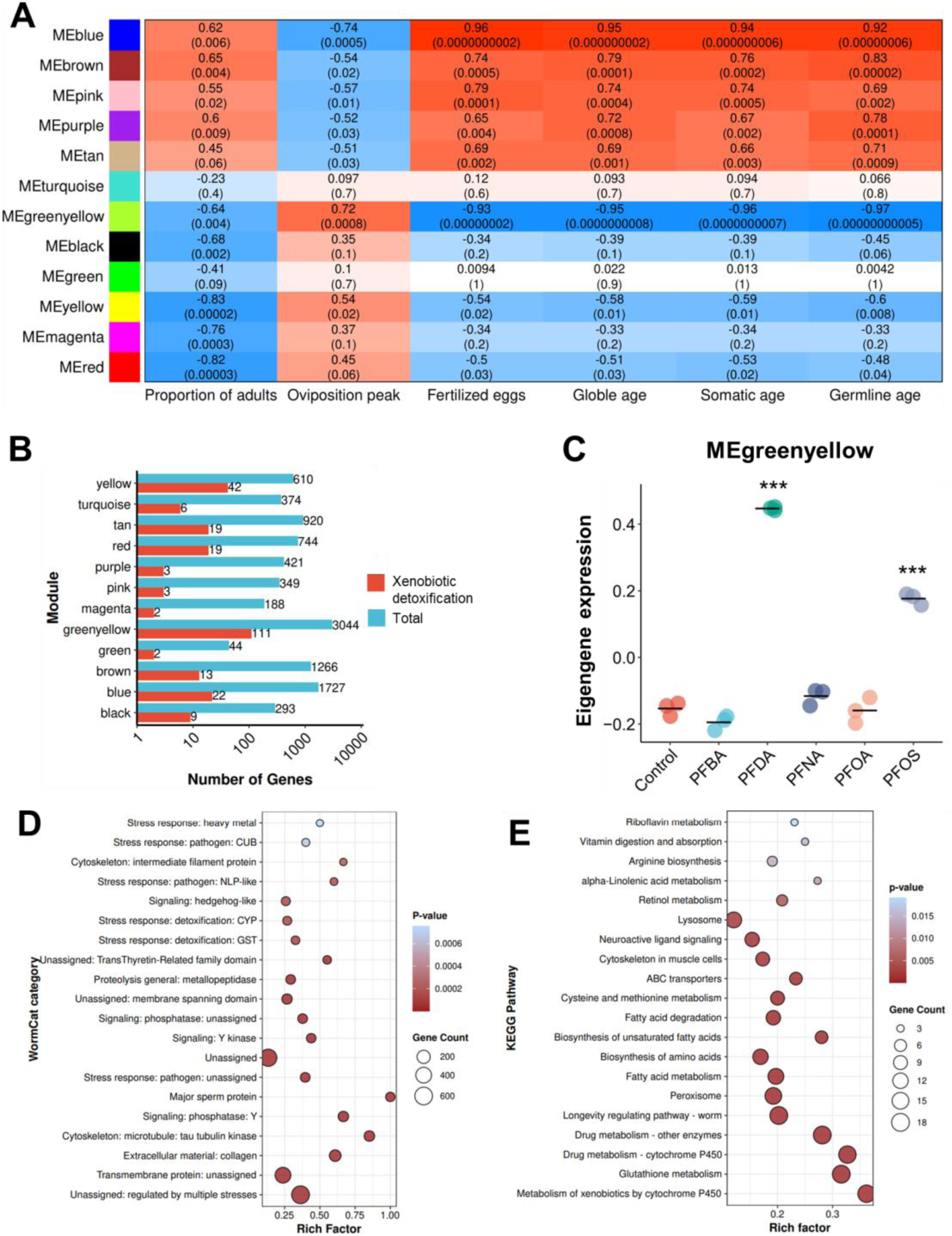
WGCNA of transcriptomic and developmental toxicity data. (**A**) Heatmap showing the correlation between identified gene modules and toxicological endpoints. Numbers in the color blocks represent correlation coefficients and corresponding p-values. (**B**) Distribution of total genes and xenobiotic detoxification genes across identified modules. (**C**) Eigengene expression of the greenyellow module across all experimental groups. Each point represents a biological replicate, with the mean line indicating the average of three replicates. (**D–E**) Enrichment analysis of genes in the greenyellow module based on (**D**) KEGG pathways and (**E**) WormCat categories. The top 20 most significantly enriched terms are shown. Statistical differences between PFAS-treated groups and the control were analyzed using one-way ANOVA with Bonferroni correction: \*\*\**p* < 0.001.

### 3.7. Alteration of transcription factors (TFs) activity induced by PFAS

Using CelEst R Shiny application, we found that PFDA and PFOS treatments resulted in quite comparable TF activity profiles. NHR-102 and NHR-85 were significantly activated (*p* < 0.001), while HLH-30 showed a pronounced suppression (*p* < 0.001; Figure 8A-B). In contrast, PFBA, PFOA, and PFNA treatments elicited distinct TF activity profiles, where PHA-4 activity was consistently decreased (*p* < 0.001; Figure 8C-F).

**Figure 8.**
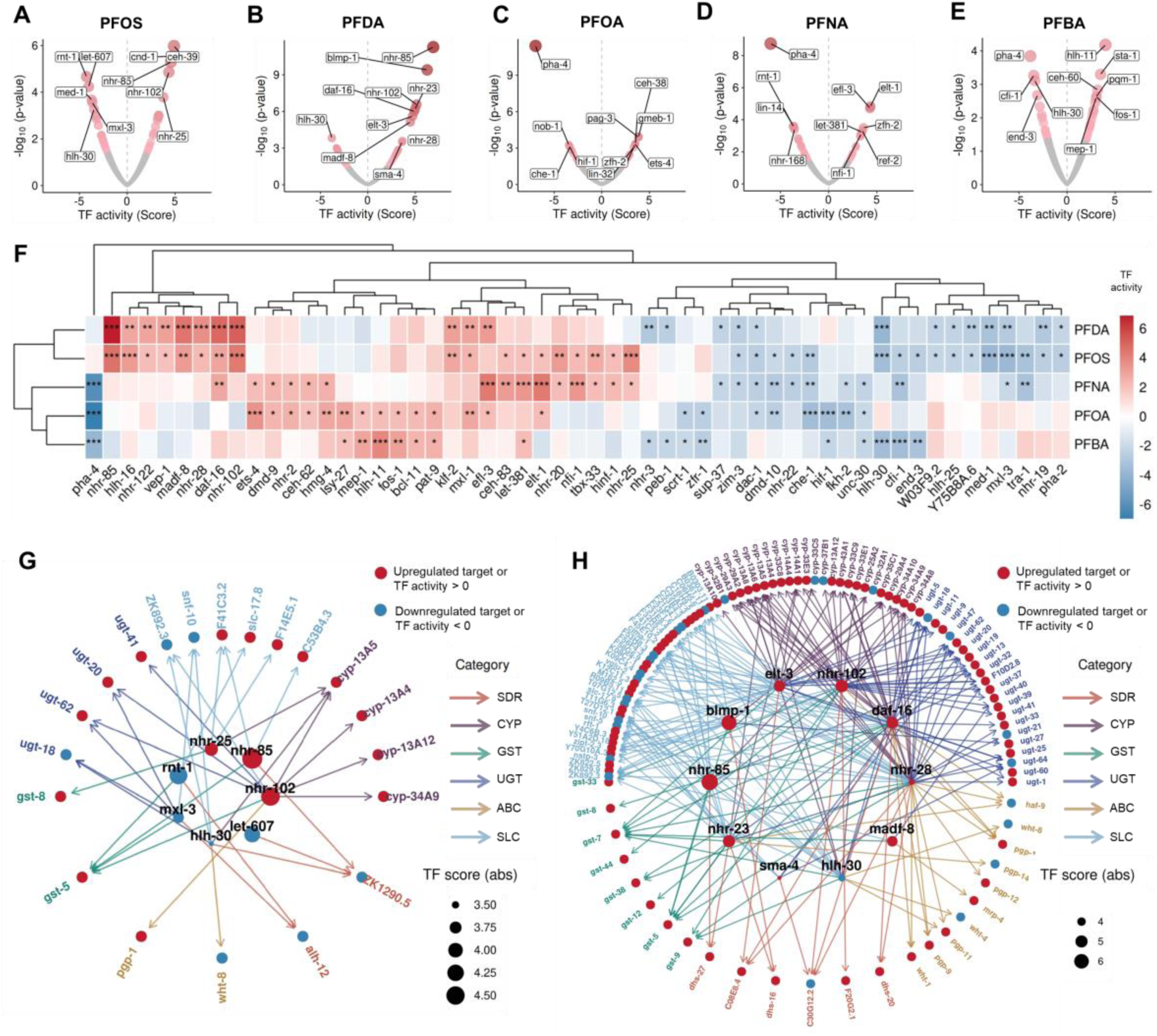
Transcription factor activity analysis across PFAS treatment groups. (**A–E**) Volcano plots displaying TF activities predicted using the CelEST shiny app. Data points are colored by statistical significance, with darker red indicating lower p-values and grey for non-significant TFs. The ten most significantly altered TFs (activated or inhibited) per group are labeled. (**F**) Heatmap of TFs significantly altered in at least two treatment groups. (**G–H**) Predicted TF regulatory networks in (**G**) PFOS and (**H**) PFDA treatment groups. Arrows link the top 10 most altered TFs to their targeted differentially expressed detoxification genes. TFs not involved in regulating xenobiotic detoxification genes are not shown in the networks. \**p* < 0.05, \*\**p* < 0.01, \*\*\**p* < 0.001.

The regulatory effects of the top 10 affected TFs in PFOS– and PFDA-treated groups on their respective age-corrected DEGs were identified using a previously constructed TF regulatory network (orthCelEsT) (Figure 8G-H) (Perez, 2025). For PFOS-treated group, seven of the top 10 TFs regulate differentially expressed xenobiotic detoxification genes (n = 20). NHR-102 targeted the largest number (n = 10) of xenobiotic detoxification genes, most of which belong to CYP family (Figure 8G). In PFDA-treated group, all top 10 altered TFs regulate differentially expressed xenobiotic detoxification genes (n = 108). NHR-28 targeted the largest number of genes (n = 44), most of which are SLC family members (Figure 8H).

### 3.8. Cross-species identification of PFAS-responsive detoxification genes linked to developmental toxicity

Cross-species analysis was conducted to determine whether transcriptomic changes related to developmental toxicity of PFAS are conserved across different species. First, overlapping genes in the WGCNA greenyellow module (MM > 0.8) and significantly upregulated xenobiotic detoxification genes were identified (Figure 9A, Table S2). This analysis yielded 24, 48, and 5 PFAS-responsive detoxification genes associated with developmental toxicity in PFOS-, PFDA-, and PFNA-treated groups, respectively (the PFBA– and PFOA-treated groups yielded no such genes). These genes were mapped to human orthologs, resulting in 29, 42, and 11 orthologs for PFOS, PFDA, and PFNA, respectively (Figure 9A, Table S2). Final filtering using data from CTD kept only those genes upregulated in at least one non-*C. elegans* species upon PFAS exposure, resulting in 14, 5, and 1 genes for PFOS, PFDA, and PFNA, respectively (Figure 9A, Table S3). These genes belong to six detoxification gene families: CYP, SDR, SLC, GST, UGT, and ABC, with most orthologs derived from humans or mice (Table S3). Notably, *CYP3A4* (ortholog of *cyp-13A2*, *cyp-13A4*, *cyp-13A5*, *cyp-13A6*, *cyp-13A7*, and *cyp-25A2*) was observed across all PFOS, PFDA, and PFNA groups. Additionally, *CBR1* (*C55A6.4*), *ABCB11* (*pgp-1*), and *GSTA5* (*gst-5*, *gst-8*) orthologs were upregulated in at least two non-*C. elegans* species under PFOS exposure (Figure 9A).

**Figure 9.**
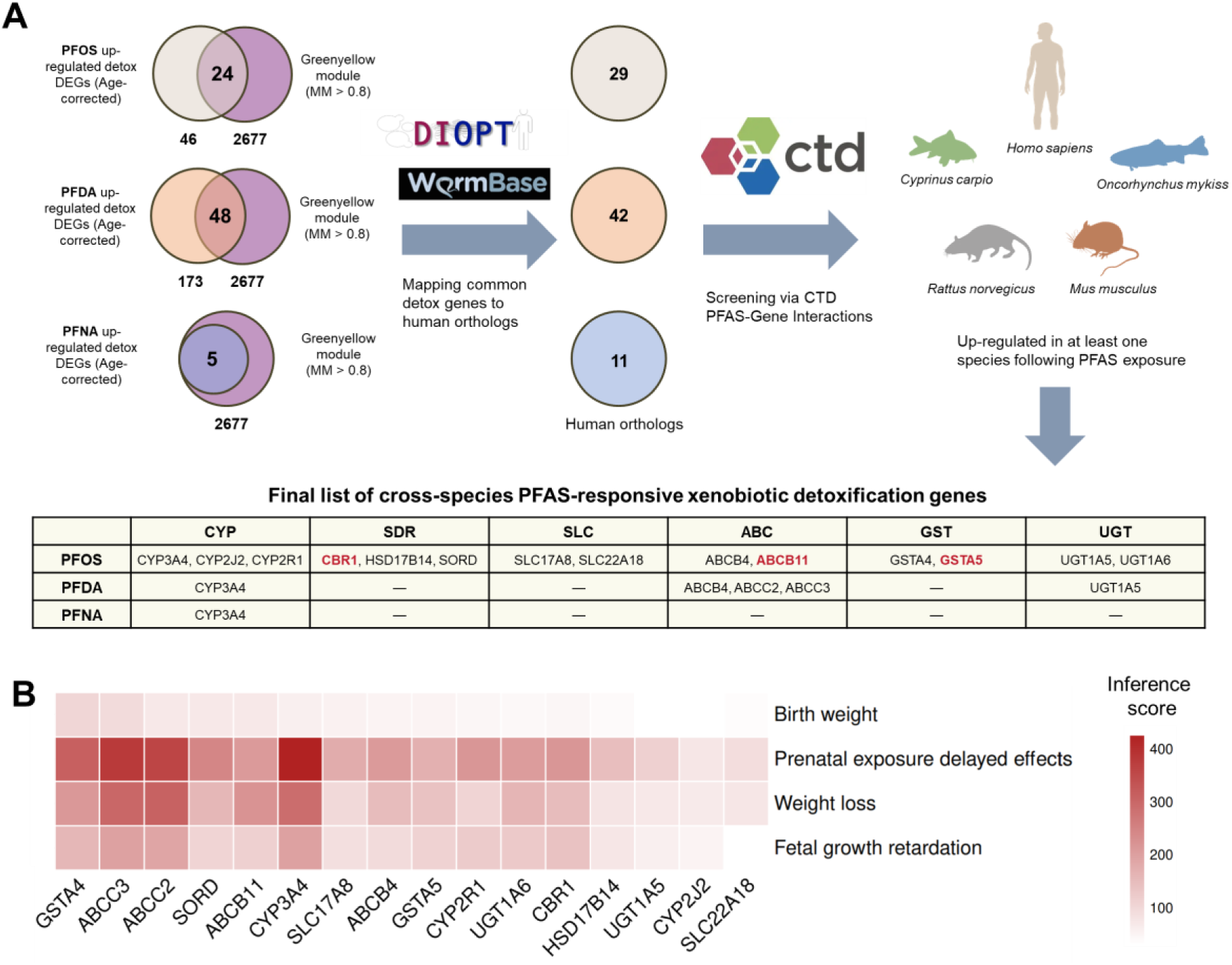
Cross-species conservation of PFAS-responsive detoxification genes and their association with developmental toxicity-related diseases. (**A**) Workflow for identifying conserved, PFAS-upregulated xenobiotic detoxification genes. Significantly upregulated, age-corrected DEGs related to xenobiotic detoxification were intersected with WGCNA Greenyellow module genes (MM > 0.8). The resulting candidates were mapped to human orthologs via WormCat (data sourced from WormBase) and DIOPT. Human homolog counts are indicated in colored circles. This gene set was filtered using CTD to retain genes upregulated in ≥1 non-*C. elegans* species (black: one; red: ≥ two). Species silhouettes: *Oncorhynchus mykiss* (by M. Michaud) and *Cyprinus carpio* (by C. Cano-Barbacil) from PhyloPic; all others from BioArt. (**B**) Disease association analysis. Genes identified in (A) were evaluated for associations with developmental toxicity-related diseases in CTD. The inference score quantifies association strength. PFBA and PFOA treatments were excluded as they induced no detoxification DEGs.

Finally, all filtered genes were analyzed for their associations with developmental toxicity-related diseases using data from CTD (Figure 9B). The term “prenatal exposure delayed effects” showed the strongest association with the entire gene set. At the gene level, *CYP3A4*, *ABCC2* (ortholog of *mrp-2*), and *ABCC3* (ortholog of *mrp-2*) exhibited the highest relevant levels with all diseases examined.

## 4. Discussion

PFAS are globally pervasive environmental contaminants having a significant bioaccumulation potential and long half-lives *in vivo* (Fenton et al., 2021). PFAS-induced developmental toxicity has been documented across diverse species, including rodents, aquatic organisms, and humans (ATSDR, 2021; Peritore et al., 2023; Truong et al., 2022). However, the molecular mechanisms underlying PFAS-mediated developmental delay remain unresolved. This study therefore evaluated the developmental delay effects of four legacy PFAS (PFDA, PFNA, PFOA, PFOS) and one emerging PFAS (PFBA) in *Caenorhabditis elegans*, and investigated their toxic mechanism using transcriptomic analysis and cross-species conservation assessment. The experimental findings are discussed in detail below.

### 4.1. Developmental inhibition induced by PFDA and PFOS

While environmentally relevant concentrations of PFNA, PFBA, and PFOA did not significantly change developmental parameters of *C. elegans*, PFOS and PFDA exhibited substantial developmental toxicity at comparable exposure levels (Figure 1B, Figure 2A-B). This structure-activity relationship aligns with previous findings in other model organisms. Specifically, long-chain PFAS generally induce more pronounced developmental toxicity than their short-chain analogues, and sulfonic acid-containing PFAS (e.g., PFOS) typically exert stronger developmental effects than carboxylic acid-containing counterparts (e.g., PFOA) (Britton et al., 2024; Kim et al., 2013). The underlying mechanism is attributed to the differences in bioaccumulation potential and biomolecular affinity among PFAS congeners, as longer chains and sulfonate groups may enhance tissue retention and interaction with cellular targets (Feng et al., 2023).

Although *in vitro* studies in other species demonstrate that PFDA impairs oocyte maturation (Domínguez et al., 2019), we did not find any notable impact of PFDA on oocyte number or brood size in *C. elegans*. The findings suggest that PFDA-induced reduction in uterine fertilized egg count is likely induced by inhibition of developmental progression, rather than disturbing gametogenesis or fertilization. PFOS exposure at concentrations exceeding 5 μM similarly suppressed development and reduced fertilized eggs in the uterus, yet showed no significant effects on oviposition timing, total brood size, or oocyte count (Figure 1B, Figure 2A-B). This can be explained by the timing of our measurements: the number of fertilized eggs in gravid adults was assessed immediately after the 72-hour exposure (starting from the L1 larval stage), whereas daily brood size was recorded starting one day after exposure termination. During this post-exposure recovery window, the subtle developmental delay induced by PFOS is likely to elicit compensatory growth. Another investigation into PFOS toxicity in *C. elegans*, using a comparable concentration (10 μM), also observed developmental suppression without finding any change in total brood size (Breton et al., 2024).

The global physiological age estimates derived from transcriptomic data were highly consistent with the developmental toxicity phenotypes observed for PFDA and PFOS (Figure 3C). Furthermore, estimates of tissue-specific ages (somatic and germline ages) revealed that PFAS-induced developmental toxicity was global (i.e., with no obvious tissue specificity) (Figure 3B). This may partially explain why PFAS induced significant developmental delay without affecting reproductive capacity. Intriguingly, we found a modest increase in physiological age of worms after PFBA treatment (Figure 3B). Previous studies have shown that PFBA treatment shortens the developmental time of *Spodoptera exigua*, and accelerates its early transition to the adult stage, suggesting that PFBA may promote the molting process of insects and nematodes (Omagamre et al., 2020).

### 4.2. PFAS exposure activates xenobiotic detoxification pathways

Separations between PFDA/PFOS groups and the control group were revealed by PCA, which mirrors the rankings of their developmental toxicity. The finding indicates that these compounds induce significant alterations in the global transcriptomic landscape (Figure 3A). However, the transcriptomic shifts are mainly caused by the differences in developmental stage (i.e., physiological age), rather than chemical perturbations (Figure S3). In *C. elegans*, even a few hours’ difference in physiological age can generate thousands of DEGs, obscuring gene expression changes directly linked to developmental toxicity (Snoek et al., 2014). Following an age-adjusted differential expression analysis, significant upregulations of xenobiotic detoxification genes—including members of the CYP, GST, UGT, SDR, SLC, and ABC transporter families— were observed in PFOS-and PFDA-exposed worms, whereas such responses were almost absent in the other three PFAS treatment groups that did not exhibit significant developmental toxicity (Figure 5-6, Figure S11).

Xenobiotic detoxification in *C. elegans* proceeds through three phases: Phase I (bioactivates the lipophilic components to provide conjugation sites for consecutive reactions, catalyzed by enzymes such as CYPs and SDRs), Phase II (conjugation with hydrophilic groups by enzymes such as GSTs and UGTs), and Phase III (efflux of metabolites by transporters including ABC and SLC families) (Hartman et al., 2021; Hoffmann and Partridge, 2015; Lindblom and Dodd, 2006). Our findings indicate coordinated activation of these detoxification systems after PFAS exposure. Among the upregulated genes, *cyp-13A4*, *cyp-13A6*, and *cyp-13A7* were the most prominently induced in PFAS-treated groups (Figure 5A-E). This expression pattern is aligned with recent findings on GenX (a PFAS alternative), indicating a conserved transcriptional response across structurally diverse PFAS (Feng et al., 2022). These CYP genes are known to be activated by a broad range of xenobiotics, including benzimidazoles (*cyp-13A6*), PCB52 (*cyp-13A6*), cadmium (*cyp-13A4*, *cyp-13A6*, *cyp-13A7*), rifampicin (*cyp-13A7*), tetrabromobisphenol A (*cyp-13A7*), and haloacetic acid disinfection by-products (*cyp-13A7*), underscoring their core roles in stress response (Larigot et al., 2022; Liu et al., 2025; Menzel et al., 2007; Stasiuk et al., 2019).

While knockdown of *cyp-13A4* has been found to increase cadmium susceptibility of worms (Cui et al., 2007), it did not alter susceptibility of worms to PFAS (Figure S7). Similar modest effects were observed in *cyp-13A6* knockdown experiments (Figure S7). The observation could be explained by non-mutually exclusive mechanisms. Specifically, given the known redundancy and broad substrate specificity of CYP families, compensatory upregulation of other detoxification genes in response to PFAS may offset the loss of *cyp-13A4* (Larigot et al., 2022; Nebert and Dalton, 2006). In addition, the exceptional stability of PFAS likely makes them recalcitrant to CYP-mediated metabolism (Hamilton et al., 2021; Pesonen and Vähäkangas, 2024). In this vein, *cyp-13A4* induction may represent a general stress response without providing significant detoxification benefits against PFAS. However, more biochemical investigations regarding *in vivo* metabolism of PFAS are required to better understand the possible roles of CYPs in response to PFAS exposure.

PPI network analysis further highlighted complex interconnections among GST, UGT, CYP, and SDR genes upon PFOS and PFDA exposure (Figure 6C-D), supporting the coordinated activation of Phase I and Phase II metabolic systems in response to PFAS stress (Du et al., 2025). In contrast, SLC and ABC transporter genes showed minor interactions with these metabolic genes. This pattern aligns with previous observations from PPI network analyses in zebrafish following xenobiotic exposure, suggesting a functional segregation between enzymatic metabolism and transmembrane efflux processes (Disner et al., 2025).

### 4.3. Transcriptional regulatory networks in PFAS-induced xenobiotic responses

TFs are pivotal regulators of development and environmental stress responses (Zhong et al., 2010). The induction of xenobiotic detoxification genes is commonly directed by specific TFs, such as nuclear hormone receptors (NHRs) (Hartman et al., 2021). However, TF activity is often modulated post-transcriptionally, and may not correlate with corresponding mRNA levels (Reinke et al., 2013). Using CelEst, we identified ten TFs showing the most significantly altered activities following PFDA and PFOS exposure, among which NHR-102, NHR-85, and HLH-30 were commonly shared (Figure 8A-F). OrthCelEsT regulatory network analysis revealed that NHR-28 and NHR-102 regulate the highest number of detoxification DEGs in PFDA and PFOS groups, respectively (Figure 8G-H). These findings indicate that PFAS activate a conserved NHR-driven transcriptional pathway for detoxification defense in *C. elegans*, reminiscing the essential role of NHR in regulating stress responses across species (Hartman et al., 2021). NHRs are ligand-dependent TFs that primarily act as transcriptional activators in modulating development, reproduction, and metabolism (Hoffmann and Partridge, 2015; Lindblom and Dodd, 2006; Zhang et al., 2025). Previous functional studies have demonstrated that *nhr-102*, *nhr-85*, and *nhr-28* are essential for overall development, vulval development, and metabolic homeostasis in *C. elegans*, respectively (Arda et al., 2010; Fuxman Bass et al., 2016; Gissendanner et al., 2004). In addition, these TFs also mediate responses to diverse xenobiotics, including dazomet, ethanol, and *P. aeruginosa* PA14 (Fuxman Bass et al., 2016; Joshi et al., 2016; Sterken et al., 2021). Our results illustrate the involvement of NHRs in the transcriptional response to PFAS-induced toxicity, suggesting a broader role for NHRs in regulating xenobiotic responses.

Transcriptional activity of HLH-30—the master regulator of autophagy and lysosomal biogenesis—was significantly reduced in PFOS, PFDA, and PFBA treatment groups (Figure 8F) (Colino-Lage et al., 2024). Although CelEst identifies TF activity without assigning a mode of regulation (e.g., activator or repressor) (Perez, 2025), the pronounced reduction in HLH-30 activity, coupled with marked downregulation of its established target genes (particularly in PFOS-treated worms), provides compelling evidence for functional inhibition of this activator (Figure 8F-H). HLH-30 regulates stress response and longevity signals in *C. elegans*, genetic loss of *hlh-30* has been shown to shorten lifespan and increase stress sensitivity of worms (Wong et al., 2023; Yu et al., 2024). The suppression of HLH-30 may inhibit cellular stress response, which in turn increases toxic effects of PFAS.

PHA-4 is a conserved TF essential for pharyngeal development and stress adaptation (Fakhouri et al., 2010; Kaiser et al., 2025). Worms lacking PHA-4 activity exhibit disrupted early pharyngeal organogenesis (Mango et al., 1994). While PHA-4 activity was significantly suppressed in PFBA, PFNA, and PFOA groups, no apparent developmental toxicity was observed (Figure 8F). It is likely that either the subtle decline of PHA-4, or the involvement of compensatory mechanisms prevents detectable morphological defects.

### 4.4. Association of PFAS-induced developmental toxicity and detoxification genes

Transcriptomic profiling revealed activation of detoxification gene regulatory networks in PFOS and PFDA groups. The functional relevance of this response was confirmed by WGCNA, which identified a “greenyellow” module that is robustly associated with developmental toxicity of PFAS (Figure 7A). Detoxification genes, particularly from CYP and GST families, were notably enriched in this module (Figure 7D-E). In addition, ME expression level of the greenyellow module across all treatment groups was highly concordant with the toxicity ranking of five PFAS (Figure 7C). Although detoxification genes generally protect worm from exogenous stress, their overexpression has been shown to disturb physiological processes through multiple mechanisms (Hartman et al., 2021). For instance, overexpression of specific GSTs can lead to developmental retardation by inducing lysosomal dysfunction, while overexpression of certain CYPs may inhibit development of worms through disrupting fatty acid metabolism (Gao et al., 2025; Min et al., 2015). Therefore, the activation of the detoxification machinery may contribute negatively to stable, CYPs-resisting chemicals in *C. elegans*. Additionally, downregulation of membrane proteins, microtubule-associated proteins, and signal transduction molecules was observed in greenyellow module (for both PFOS and PFDA treatment groups) (Figure 7D-E, Figure S8), suggesting the involvement of cytoskeletal structure/signal transduction pathways in modulating PFAS-induced developmental toxicity.

Dozens of PFAS-induced detoxification genes in *C. elegans* have been reported in other species, underscoring the conserved role of these genes in mediating PFAS-induced transcriptomic responses (Table S2-3). Notably, CYP3A4 and its orthologs are shared stress-responsive genes across PFDA, PFOS, and PFNA treatments in both human cells and *C. elegans*, suggesting its potential as a cross-species biomarker of PFAS toxicity (Figure 9A, Table S2-3). The transcriptional signature of PFOS exposure indicates a comprehensive xenobiotic response, with consistent upregulation of six major detoxification gene families was observed (Figure 9B). Among these genes, the orthologs of CBR1 (SDR family), GSTA5 (GST family), and ABCB11 (ABC transporter family) were upregulated in at least two non-*C. elegans* species, indicating a remarkably cross-species conservation of these genes in response to PFAS. While most studies show stable upregulation of these genes after PFAS exposure, contradictory findings have been reported, indicating that certain dosing thresholds could be required for triggering PFAS-induced detoxification responses (Franco et al., 2020). Additionally, the limited toxicological data of PFNA and PFDA may introduce biases in database-derived analysis, underscoring the need for investigations of cross-species detoxification mechanisms covering a wide range of PFAS substances.

Disease association analysis showed that CYP3A4 exhibited the strongest association with development-related health issues, particularly “Prenatal Exposure Delayed Effects” (Figure 9B). Although CYP3A4 overexpression can enhance cellular detoxification capacity, certain xenobiotics may generate more toxic metabolites during biotransformation, leading to significant developmental disturbances (Hakkola et al., 1998). Furthermore, since CYP3A4 extensively metabolizes endogenous substrates such as vitamin D and steroid hormones, its overexpression can disrupt these critical processes, leading to specific developmental disorders including rickets and impaired organ development (Klyushova et al., 2022; Roizen et al., 2018; Yu et al., 2005).

Apart from CYP3A4, significant associations between transporters ABCC2/ABCC3 and developmental toxicity-related diseases were also observed (Figure 9B). While efflux activation usually exerts a protective action against xenobiotics, ABCC2/ABCC3 overexpression can deplete intracellular glutathione (GSH) via efflux, potentially inhibiting cell growth (Brechbuhl et al., 2012; Park, 2016; Wang et al., 2024). The findings suggest double-edged roles of detoxification genes in mediating PFAS-induced developmental toxicity, which warrants further investigations.

## 5. Conclusion

In summary, our results reveal that PFAS exert developmental toxicity in *C. elegans* through a mechanism closely linked to the activation of detoxification genes. The conservation of these transcriptional responses across species highlights detoxification enzymes, particularly CYP3A4 and its orthologs, as key candidates for further mechanistic investigation and biomarker development. While this study establishes strong associations between developmental toxicity of PFAS and specific genes, future genetic manipulation studies are essential to understand their exact roles within the adverse outcome pathway (AOP) framework.

## Data and code availability

The raw bulk RNA-seq data have been deposited in the Genome Sequence Archive (GSA) under the accession number CRA031423. The scripts used to generate the standard and age-corrected DEGs, along with the corresponding count files, are available at: https://github.com/ZhenxiaoCao/Transcriptomic-insights-into-developmental-toxicity-of-PFAS-in-Caenorhabditis-elegans.git

## Acknowledgements

We thank the Caenorhabditis Genetics Center for providing *C. elegans* strains. We also extend our gratitude to Dr. Mirko Francesconi and Dr. Romain Bulteau (Université Claude Bernard Lyon 1) for their technical assistance in using RAPToR.

