## Supplementary material for "Transcriptomic insights into developmental toxicity of per- and poly-fluoroalkyl substances (PFAS) in *Caenorhabditis elegans*: The potential key role of xenobiotic detoxification pathway": Text S1, Figure S1-S13, Table S1-S3

**Table of Contents**

**Supplementary Text**

Text S1. Details of methods

**Supplementary Figures**

Figure S1. Total brood size of *C. elegans* following 60-hour exposure to PFAS

Figure S2. Oocyte counts in *C. elegans* after 72-hour exposure to PFAS

Figure S3. Correlation between observed transcriptomic changes and shifts attributable to developmental age

Figure S4. Overlap of age-corrected DEGs across PFAS treatments

Figure S5. Analysis of standard DEGs across PFAS treatments

Figure S6. Validation of key CYP genes expression by RT-qPCR

Figure S7. RNAi knockdown of CYP genes partially rescues PFAS toxicity

Figure S8. Functional enrichment analysis of downregulated age-corrected DEGs

Figure S9. WormCat enrichment analysis of standard DEGs

Figure S10. KEGG pathway enrichment analysis of standard DEGs

Figure S11. Quantification of age-corrected, differentially expressed detoxification genes across PFAS treatments

Figure S12. Selection of the soft‑threshold power for WGCNA

Figure S13. Eigengene expression profiles of WGCNA modules across treatment groups

**Supplementary Tables**

Table S1. Primers used for RT-qPCR

Table S2. Intersection between upregulated detoxification-related DEGs (age-corrected) in PFAS treatment groups and the Greenyellow module (MM ≥ 0.8), along with WormCat functional annotations and human orthologs

Table S3. Cross-species PFAS-responsive xenobiotic detoxification genes and corresponding literature evidence

**Text S1. Details of methods**

*Daily brood size*

Following PFAS exposure, individual young adult worms were transferred to fresh NGM plates every 24 hours. After two days of incubation at 20 ℃, eggs and hatched larvae on each plate were counted under a stereomicroscope (Motic, Xiamen, China), and these counts represented the daily brood size. At least five worms were analyzed per group. The brood size–time relationship was modeled using polynomial curve fitting to estimate the time of oviposition peak, which was subsequently used for correlation analysis.

*RNAi treatment*

For RNAi treatment, a single colony of *E. coli* HT115 bacteria expressing dsRNA targeting the gene of interest was inoculated into LB broth supplemented with ampicillin (100 mg/L) and tetracycline (5 mg/L). The culture was then grown overnight at 37 °C with shaking at 200 rpm. Isopropyl β-D-1-thiogalactopyranoside (IPTG, 0.5 mM) was then added to the culture, which was further incubated for 4 hours to induce dsRNA expression. To increase bacterial density, the bacterial culture was concentrated 50-fold by centrifugation (5000 × g for 15 minutes) and resuspended in M9 buffer. The concentrated bacterial suspension was then used to feed L1-stage worms, which were simultaneously exposed to PFAS. HT115 bacteria harboring the empty L4440 vector were used as the control group.

*WGCNA*

WGCNA was performed to identify associations between transcriptomic alterations and developmental toxicity traits. Among the six analyzed traits (global age, somatic age, germline age, proportion of adults, oviposition peak, and fertilized egg count), three (global age, somatic age, and germline age) were directly estimated from the transcriptome data, enabling strict one-to-one matching with the RNA-seq samples. However, due to an insufficient number of individuals from the sequenced nematodes for phenotypic assays, the remaining three traits (proportion of adults, oviposition peak, and number of fertilized eggs) were measured using a separate batch of nematodes in independent experiments. Consequently, these three traits could not be paired on a per-sample basis with the transcriptome data.

To minimize the potential impact of this limitation, we used the mean values of proportion of adults, oviposition peak, and fertilized egg count across biological replicates in the WGCNA correlation analysis. While this approach does not provide sample-level pairing, it allows for an approximate integration of these traits with the transcriptome modules and reduces potential bias.

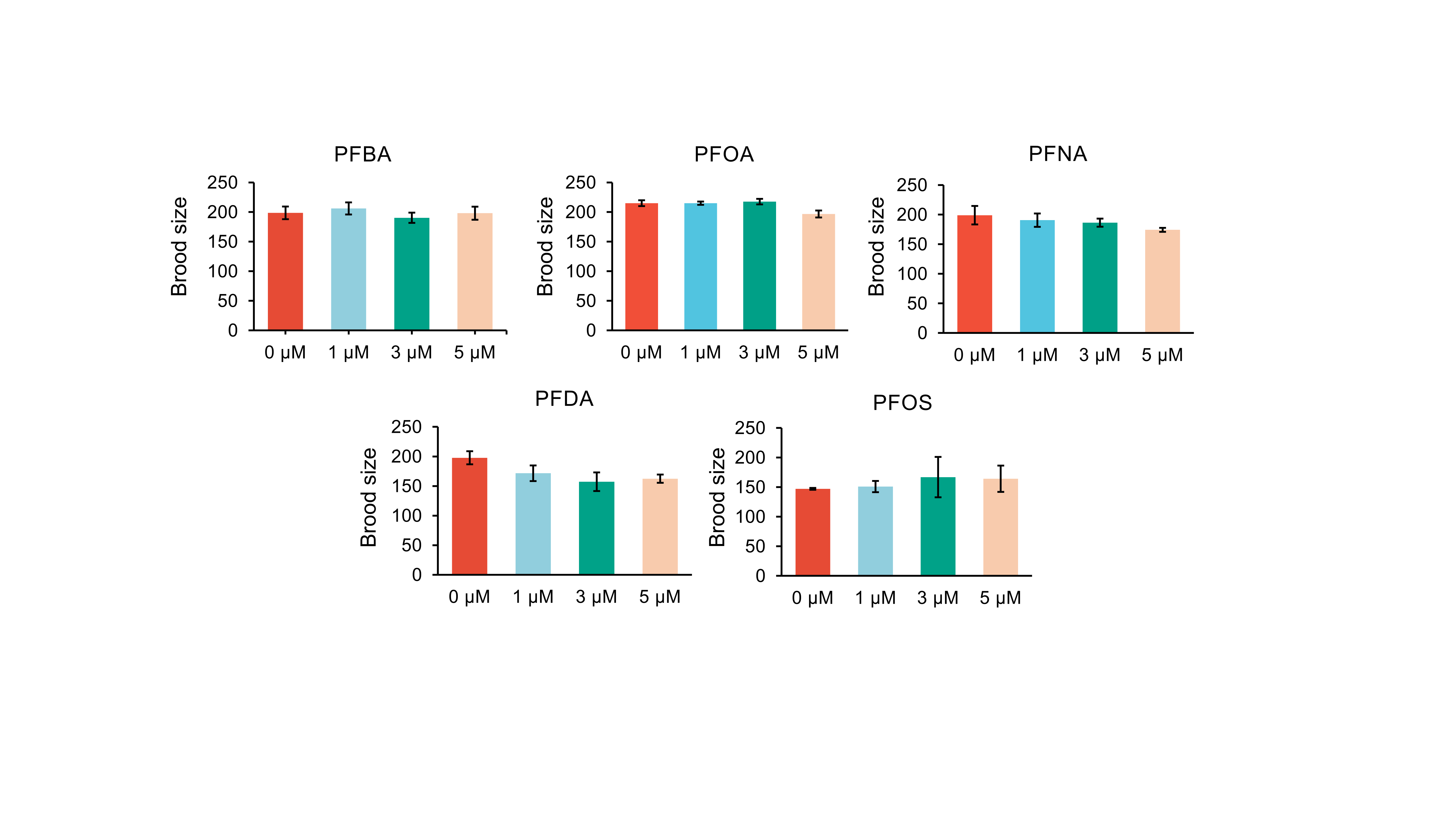

**Figure S1.** Total brood size of *C. elegans* following 60-hour exposure to PFAS. Data are presented as mean ± SEM from at least three independent biological replicates.

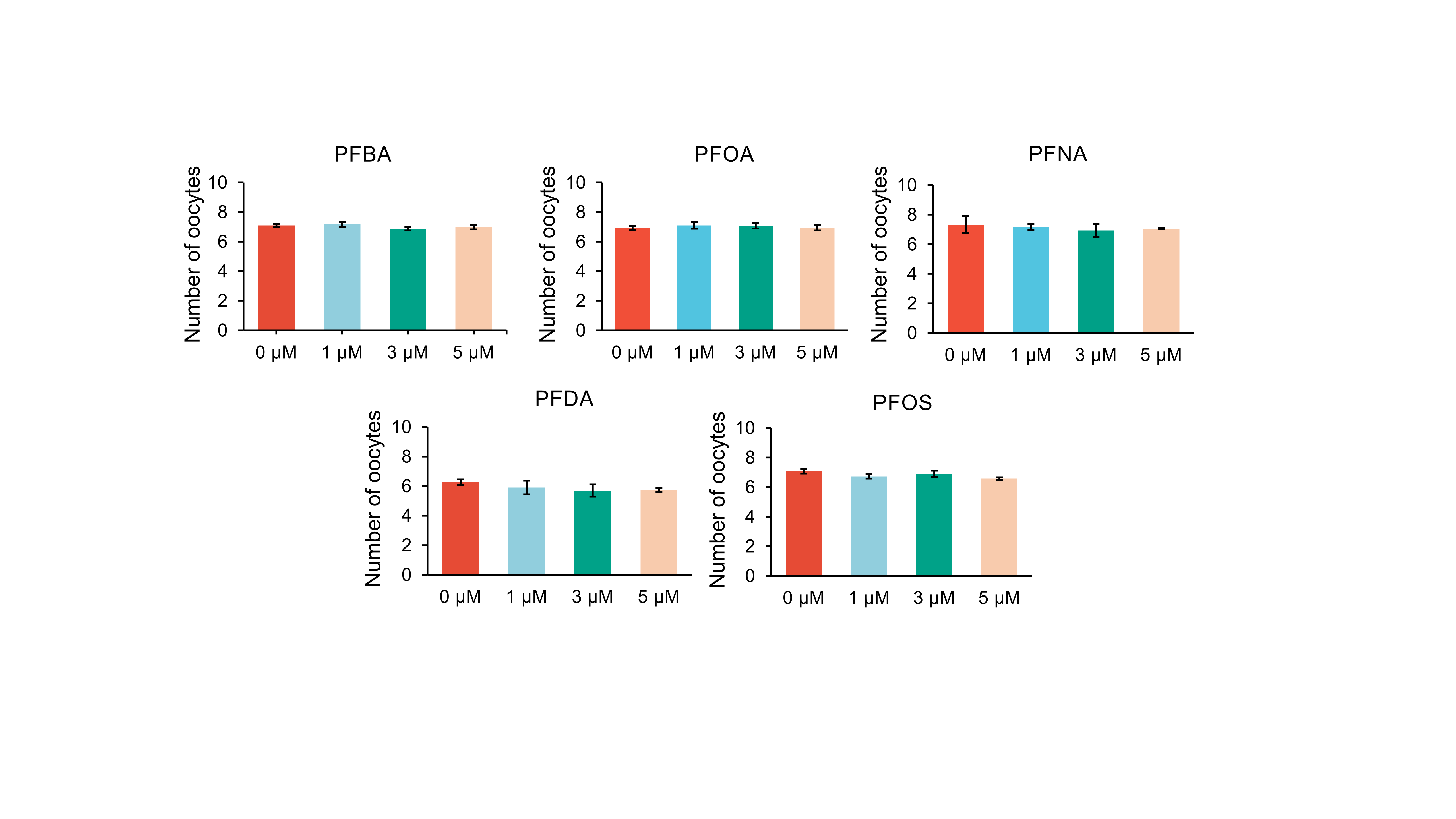

**Figure S2**. Oocyte counts in *C. elegans* after 72-hour exposure to PFAS. Data are presented as mean ± SEM from at least three independent biological replicates.

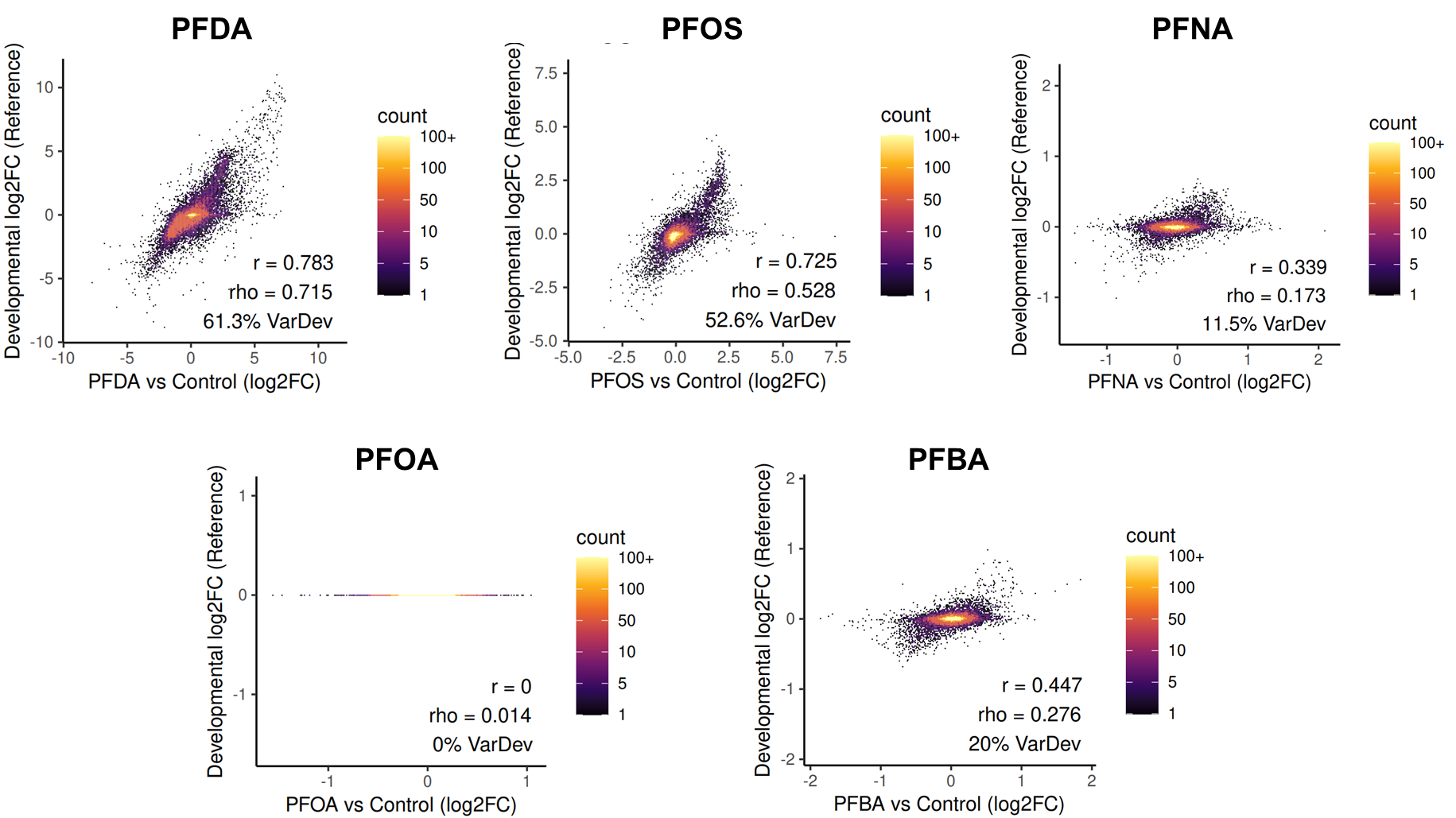

**Figure S3.** Correlation between observed transcriptomic changes and shifts attributable to developmental age. Scatter plots comparing the observed log₂ fold changes (log₂FC) in gene expression (PFAS vs. control) against the log₂FC predicted from developmental time differences alone. The Pearson (r) and Spearman (rho) correlation coefficients, and variance explained by development (VarDev, calculated as r²) are indicated for each treatment.

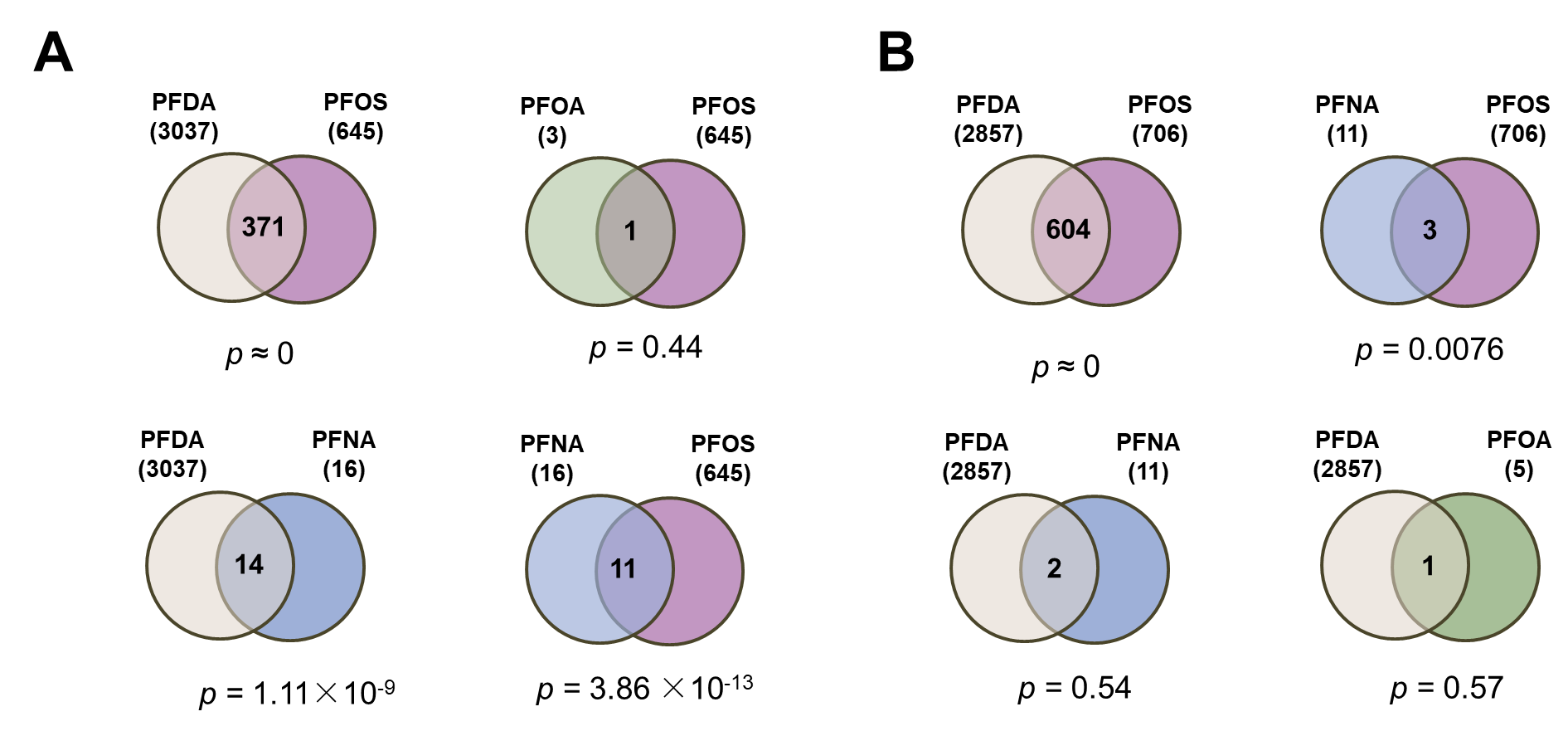

**Figure S4**. Overlap of age-corrected DEGs across PFAS treatments. Venn diagrams depict the pairwise overlaps of (A) upregulated and (B) downregulated age-corrected DEGs among the different PFAS treatment groups. The statistical significance of the overlaps was assessed using the hypergeometric test.

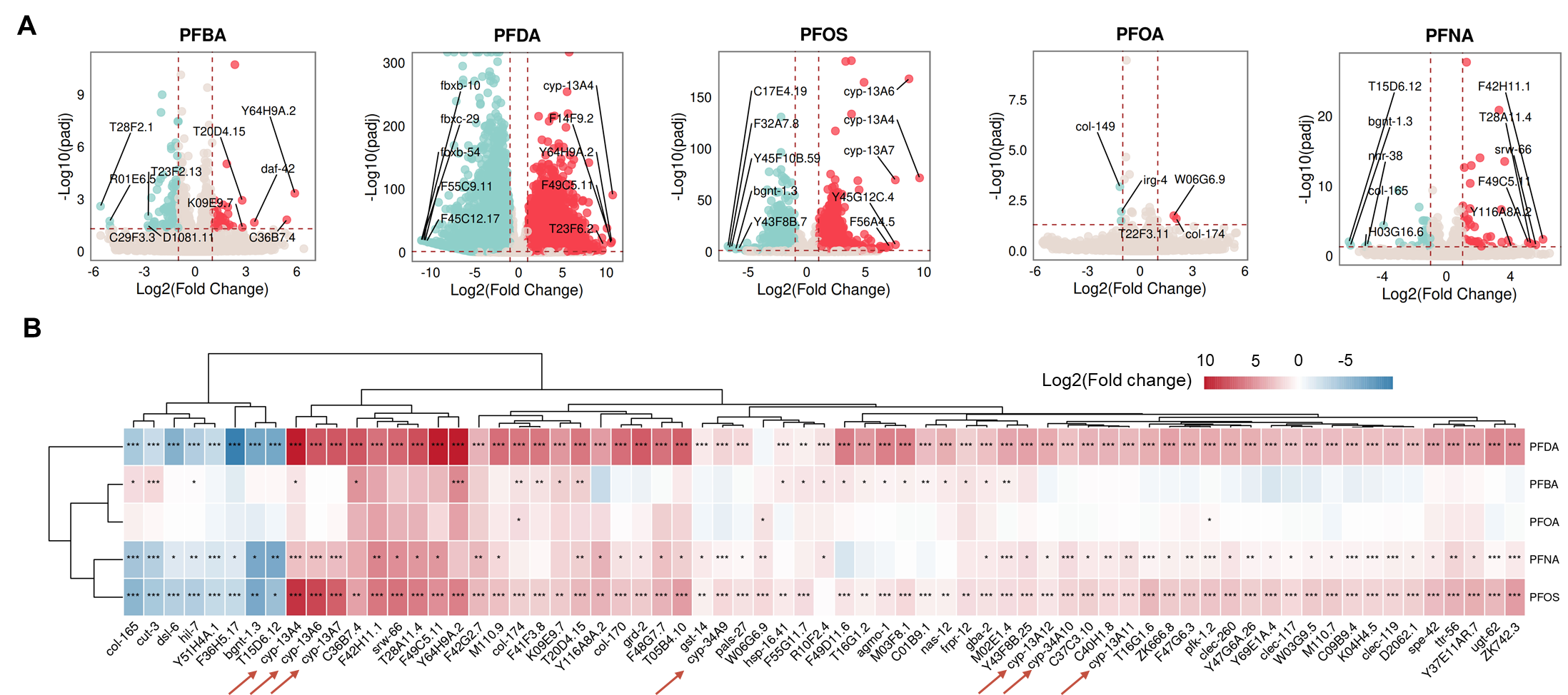

**Figure S5.** Analysis of standard DEGs across PFAS treatments. (A) Volcano plots of standard DEGs for each PFAS treatment. The top five most significantly upregulated and downregulated genes in each group are labeled. (B) Heatmap displaying DEGs that were consistently and significantly dysregulated in at least three treatment groups. Arrows highlight CYP genes. * adjusted *p* < 0.05, ** adjusted *p* < 0.01, *** adjusted *p* < 0.001.

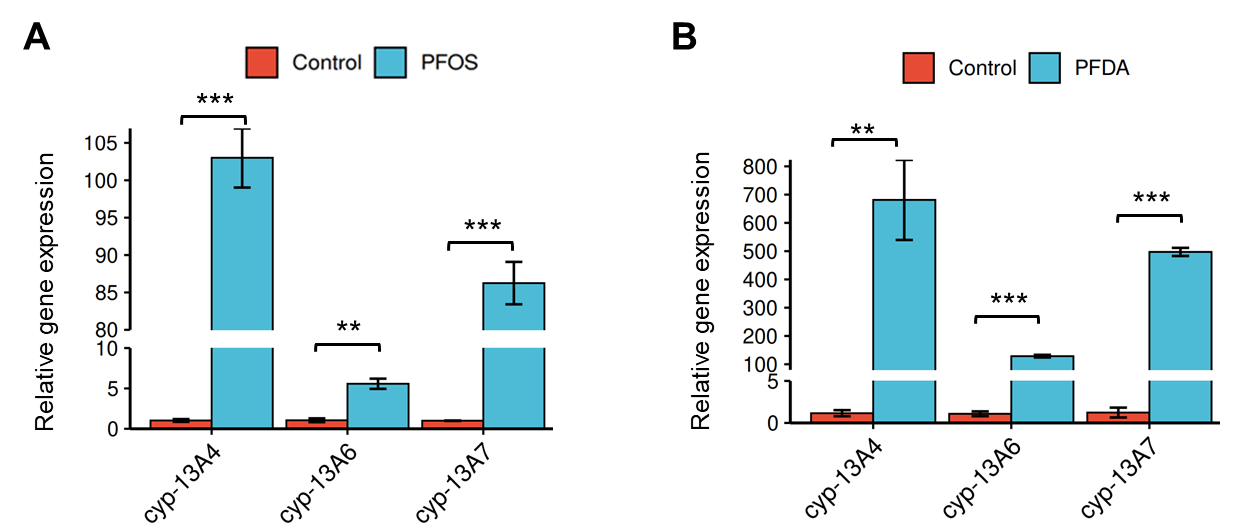

**Figure S6.** Validation of key CYP genes expression by RT-qPCR. Relative mRNA expression levels of *cyp-13A4*, *cyp-13A6*, and *cyp-13A7* in worms exposed to (A) PFOS or (B) PFDA. Data were normalized to the reference gene *tba-1* and are presented as mean ± SEM from at least three independent experiments. Statistical significance was determined by two-tailed Student's t-test (***p* < 0.01, ****p* < 0.001).

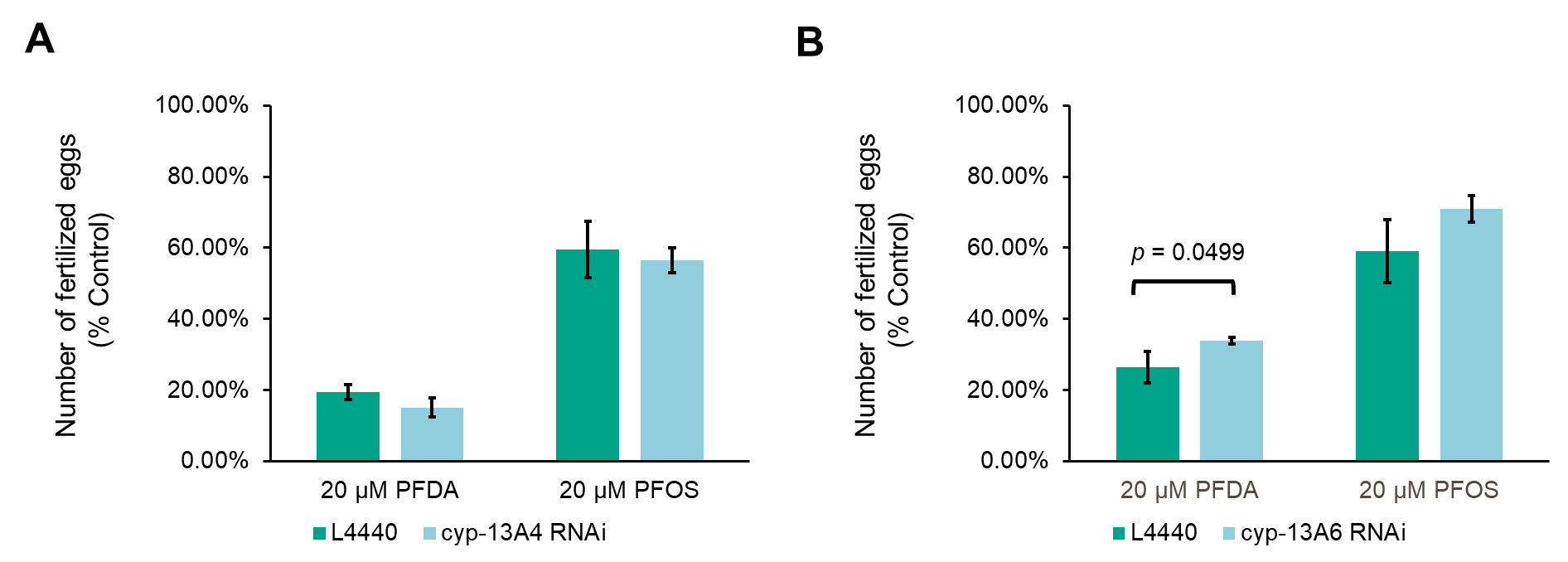

**Figure S7.** RNAi knockdown of CYP genes partially rescues PFAS toxicity. The number of fertilized eggs was quantified in worms fed with (A) *cyp-13A4* or (B) *cyp-13A6* RNAi bacteria following 72-hour exposure to PFAS, starting from the L1 larval stage. The empty vector L4440 was used as control. *p*-values were calculated by two-tailed Student's t-test.

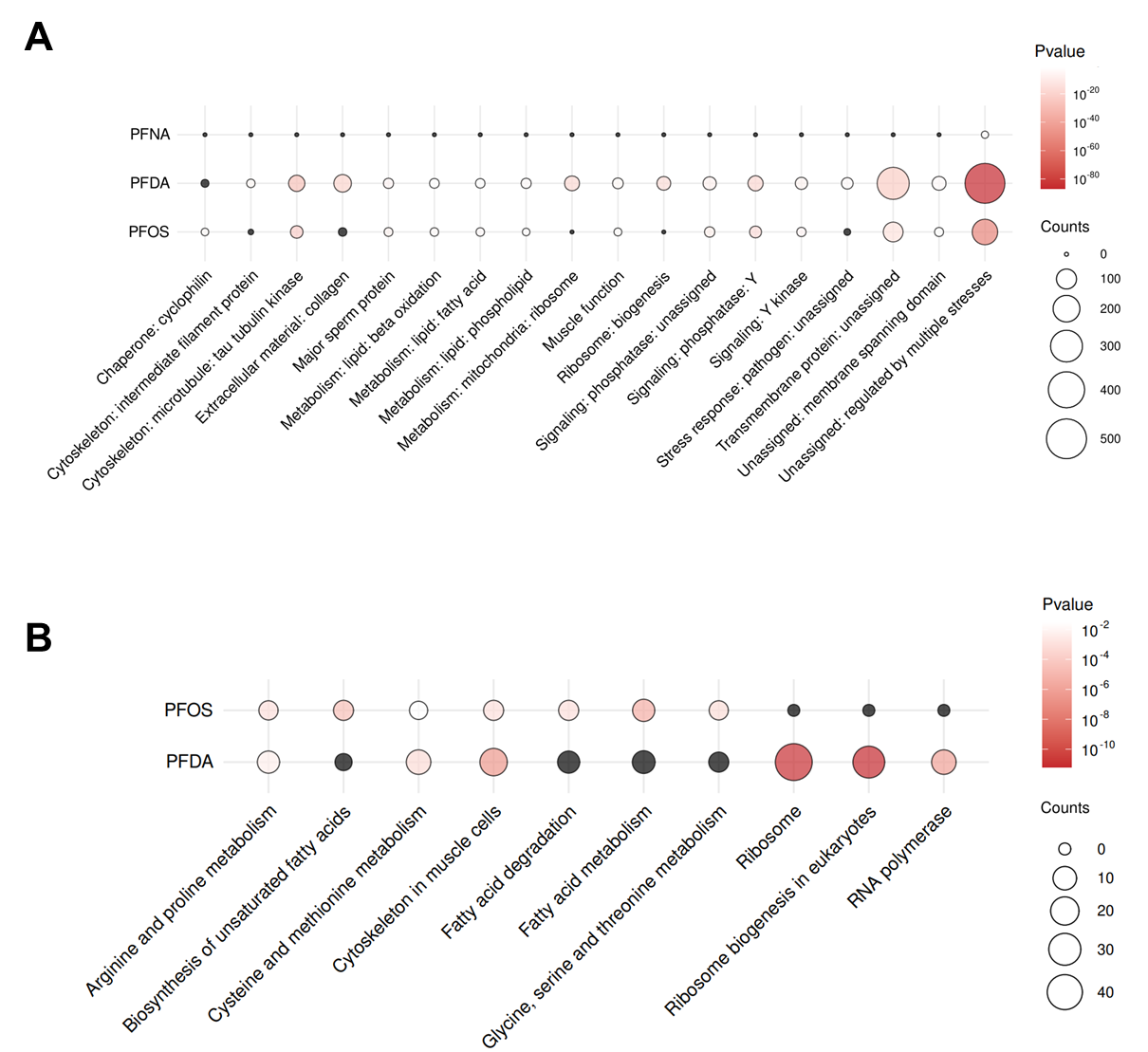

**Figure S8.** Functional enrichment analysis of downregulated age-corrected DEGs. Enriched terms from (A) WormCat and (B) KEGG pathway analyses are shown. Only terms with a *p*-value < 0.001 in at least one treatment group are displayed. Black bubbles represent non-significant terms.

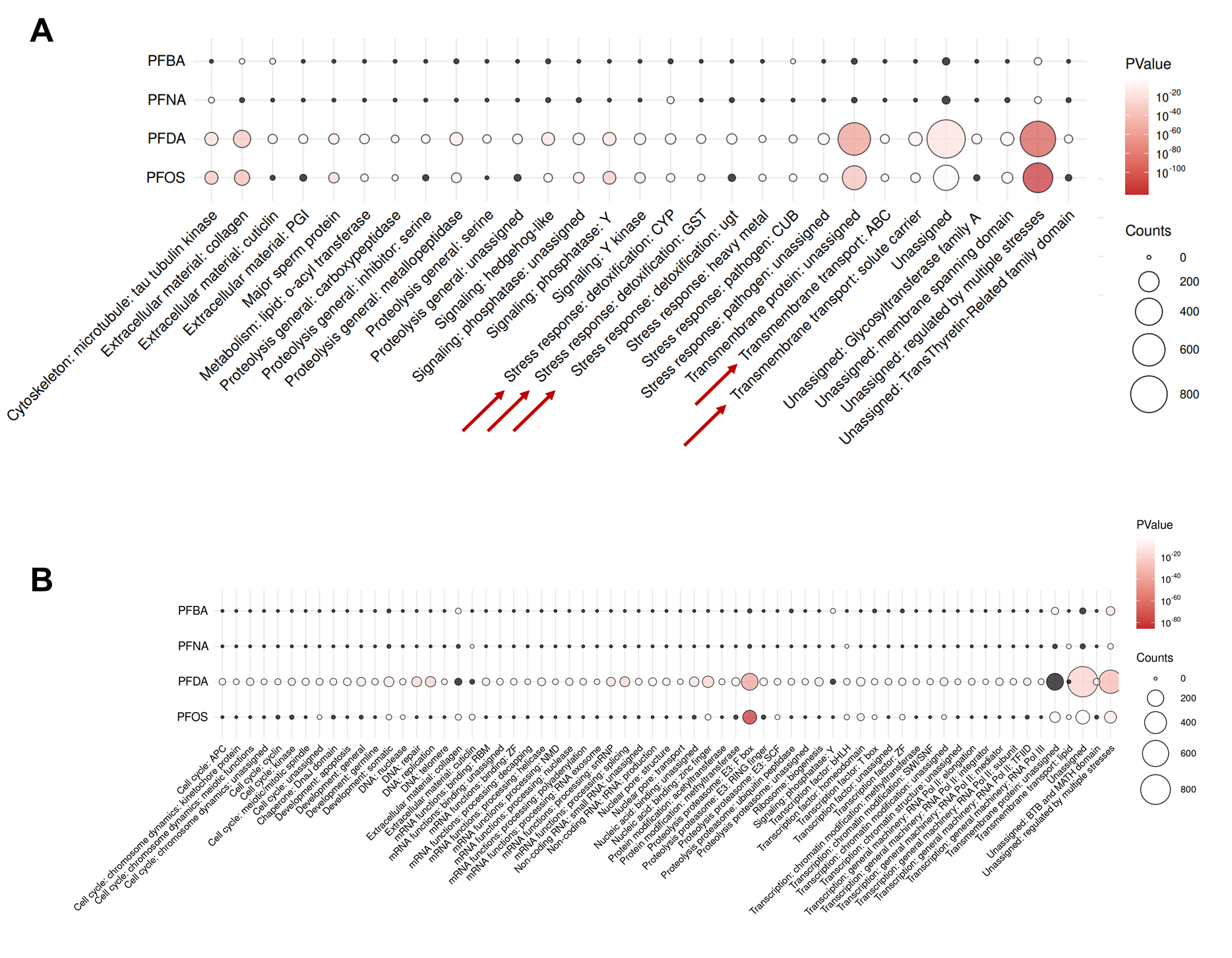

**Figure S9.** WormCat enrichment analysis of standard DEGs. Enrichment results for (A) upregulated and (B) downregulated standard DEGs. Only terms with a *p*-value < 0.001 in at least one treatment group are displayed. Black bubbles represent non-significant terms. Arrows indicate xenobiotic detoxification categories.

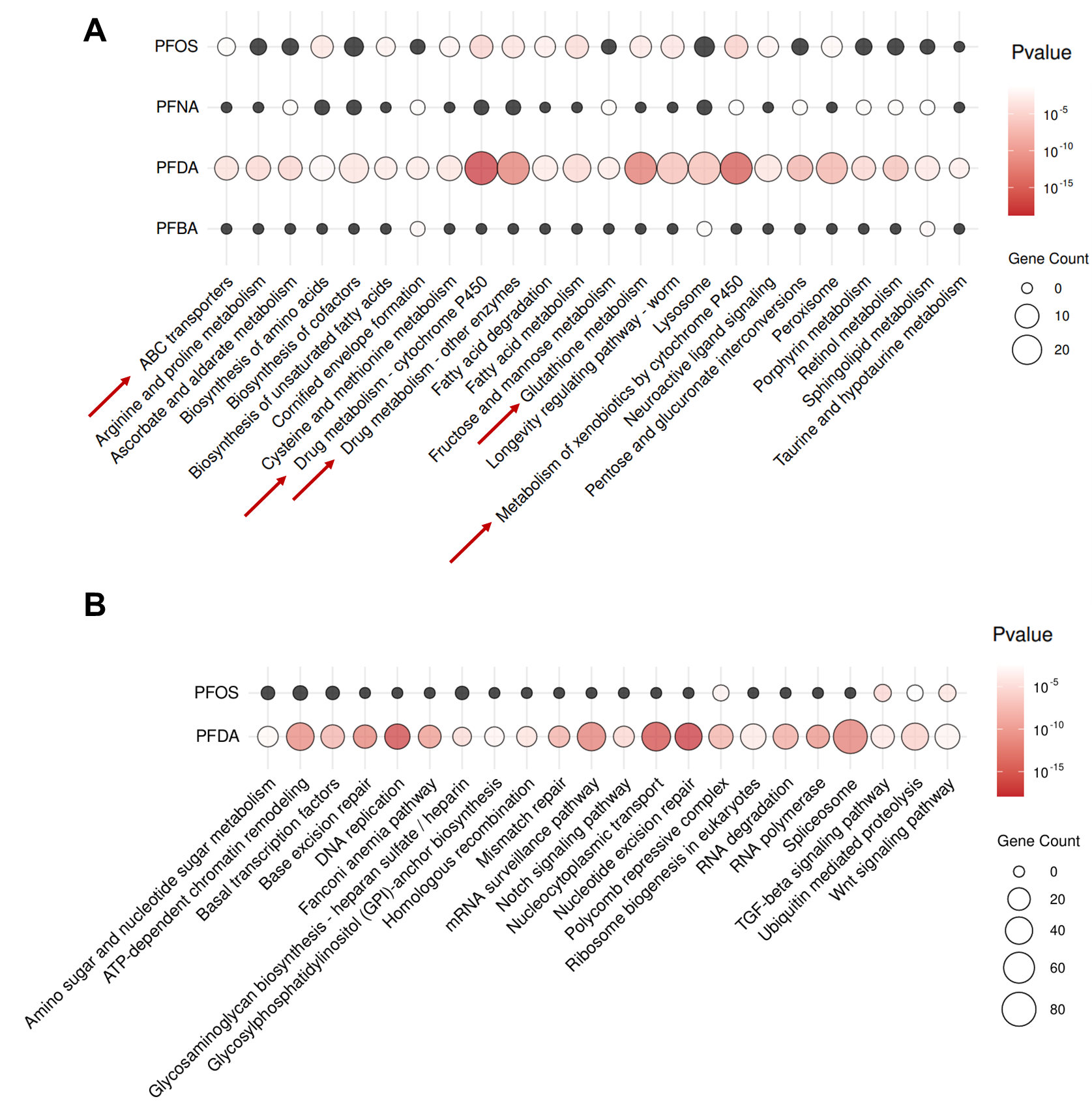

**Figure S10.** KEGG pathway enrichment analysis of standard DEGs. Enrichment results for (A) upregulated and (B) downregulated standard DEGs. Only terms with a *p*-value < 0.001 in at least one treatment group are displayed. Black bubbles represent non-significant terms. Arrows indicate xenobiotic detoxification categories.

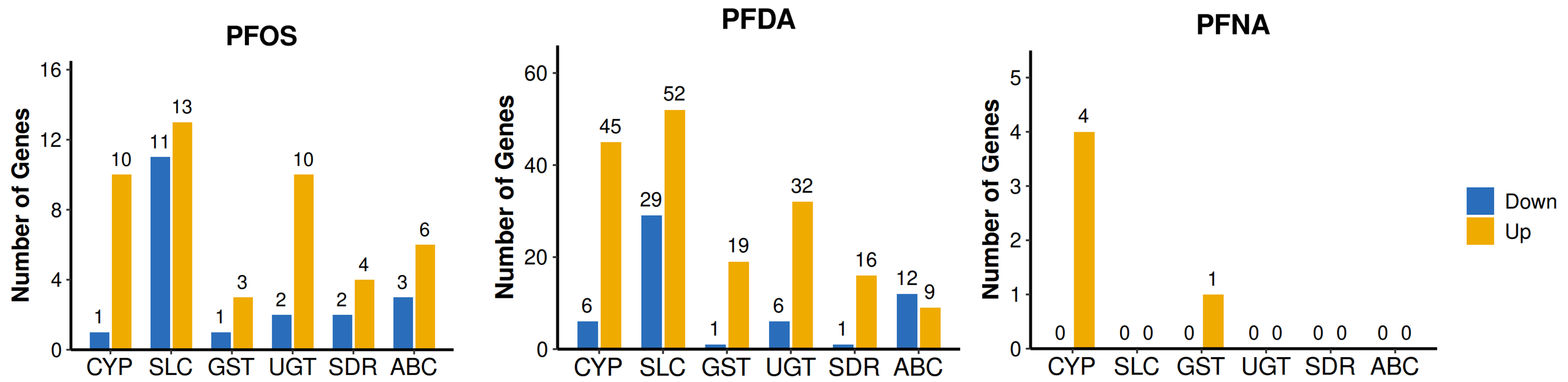

**Figure S11.** Quantification of age-corrected, differentially expressed detoxification genes across PFAS treatments.

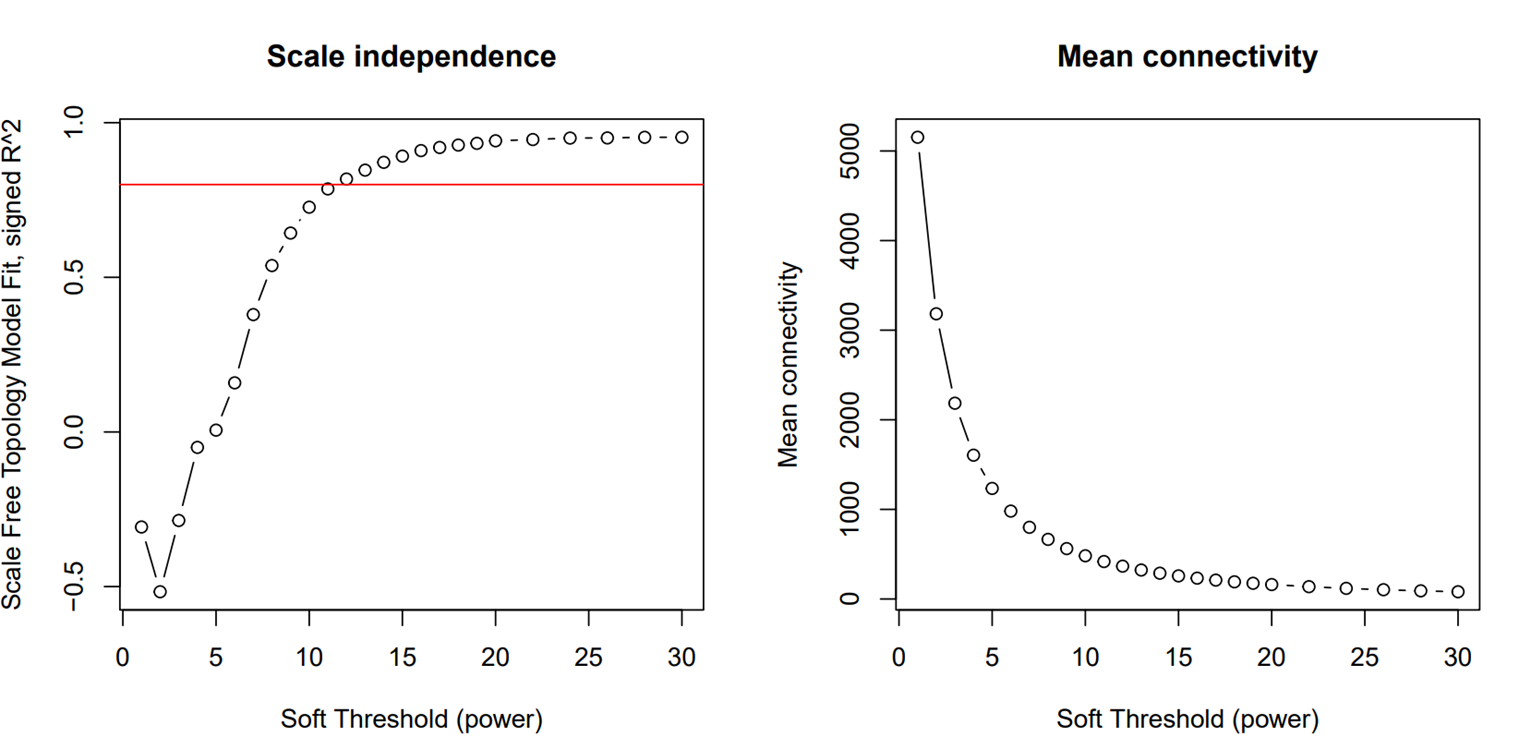

**Figure S12.** Selection of the soft‑threshold power for WGCNA. A power of 12 was selected as the lowest value achieving a scale‑free topology fit index (R²) ≥ 0.8 while retaining relatively high mean connectivity.

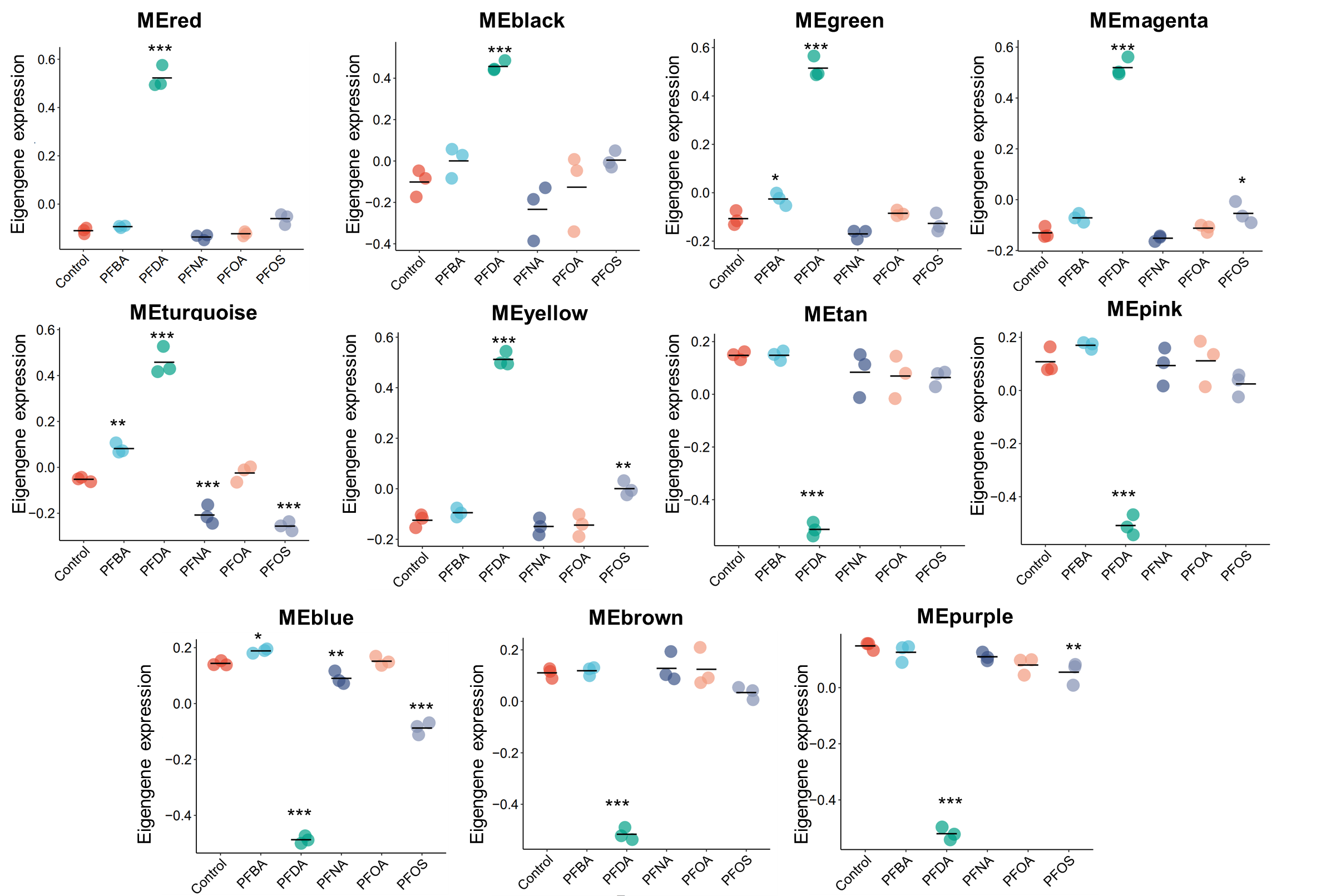

**Figure S13.** Eigengene expression profiles of WGCNA modules across treatment groups. Each point represents a biological replicate, with the mean line indicating the average of three replicates. Statistical significance between PFAS-treated groups and the control was determined by one-way ANOVA with Bonferroni correction. **p* < 0.05, ***p* < 0.01, ****p* < 0.001.

**Table S1.** Primers used for RT-qPCR.

| Gene name | Sequence of the primer (5’-3’) | Reference |
| --- | --- | --- |
| *cyp-13A4* | F: GGTCAAACCGAATCGCTGATG R: TTGGCGCATCAAAGTTCCTG | Thapa I, Vahrenkamp R, Witus SR, et al. Nucleic Acids Res. 2023;51(5):2108-2116. |
| *cyp-13A6* | F: CGTGATCTCCAGAGTTGCCA  R: AACATCCAGATCTGCCAGCG |  |
| *cyp-13A7* | F: TGTTAACGAGTTCAAGCCGGA  R: ATTCTTGGTCCCATGCCGAA |  |
| *tba-1* | F: TCAACACTGCCATCGCCGCC  R: TCCAAGCGAGACCAGGCTTCAG |  |

**Table S2.** Intersection between upregulated detoxification-related DEGs (age-corrected) in PFAS treatment groups and the Greenyellow module (MM ≥ 0.8), along with WormCat functional annotations and human orthologs.

| PFAS Treatment | Category (WormCat) | Gene Name | Human Homologs |
| --- | --- | --- | --- |
| PFDA | Metabolism: short chain dehydrogenase | *C55A6.4* | CBR1 |
| PFDA | Metabolism: short chain dehydrogenase | *C55A6.6* | CBR3 |
| PFDA | Metabolism: short chain dehydrogenase | *C55A6.7* | CBR1 |
| PFDA | Metabolism: short chain dehydrogenase | *D2063.1* | SORD |
| PFDA | Metabolism: short chain dehydrogenase | *dhs-20* | HSD17B6, DHRS9, RDH5 |
| PFDA | Metabolism: short chain dehydrogenase | *F20G2.1* | CBR3 |
| PFDA | Stress response: detoxification: CYP | *cyp-13A11* | CYP3A7-CYP3A51P |
| PFDA | Stress response: detoxification: CYP | *cyp-13A12* | CYP3A7-CYP3A51P |
| PFDA | Stress response: detoxification: CYP | *cyp-13A2* | TBXAS1, CYP3A7, CYP3A5, CYP3A4 |
| PFDA | Stress response: detoxification: CYP | *cyp-13A4* | TBXAS1, CYP3A7, CYP3A5, CYP3A4 |
| PFDA | Stress response: detoxification: CYP | *cyp-13A5* | TBXAS1, CYP3A7, CYP3A5, CYP3A4 |
| PFDA | Stress response: detoxification: CYP | *cyp-13A6* | TBXAS1, CYP3A7, CYP3A5, CYP3A4 |
| PFDA | Stress response: detoxification: CYP | *cyp-13A7* | TBXAS1, CYP3A7, CYP3A5, CYP3A4 |
| PFDA | Stress response: detoxification: CYP | *cyp-25A2* | TBXAS1, CYP3A43, CYP3A7, CYP3A4 |
| PFDA | Stress response: detoxification: CYP | *cyp-33C1* | CYP2J2, CYP2R1, CYP2U1 |
| PFDA | Stress response: detoxification: CYP | *cyp-33C8* | CYP2J2, CYP2R1, CYP2U1 |
| PFDA | Stress response: detoxification: CYP | *cyp-33C9* | CYP2J2, CYP2R1, CYP2U1 |
| PFDA | Stress response: detoxification: CYP | *cyp-33E3* | CYP21A2, CYP2U1, CYP2J2, CYP2R1 |
| PFDA | Stress response: detoxification: CYP | *cyp-34A10* | CYP21A2, CYP2U1, CYP2J2, CYP2R1 |
| PFDA | Stress response: detoxification: CYP | *cyp-34A8* | CYP21A2, CYP2U1, CYP2J2, CYP2R1 |
| PFDA | Stress response: detoxification: CYP | *cyp-34A9* | CYP21A2, CYP2U1, CYP2J2, CYP2R1 |
| PFDA | Stress response: detoxification: CYP | *cyp-43A1* | CYP3A43, CYP3A5 |
| PFDA | Stress response: detoxification: GST | *gst-19* | GSTA4, GSTA5, HPGDS |
| PFDA | Stress response: detoxification: GST | *gst-29* | GSTA4, GSTA5, HPGDS |
| PFDA | Stress response: detoxification: GST | *gst-33* | GSTA4, GSTA5, HPGDS |
| PFDA | Stress response: detoxification: GST | *gst-38* | GSTA4, GSTA5, HPGDS |
| PFDA | Stress response: detoxification: GST | *gst-5* | GSTA4, GSTA5, HPGDS |
| PFDA | Stress response: detoxification: GST | *gst-7* | GSTA4, GSTA5, HPGDS |
| PFDA | Stress response: detoxification: GST | *gst-8* | GSTA4, GSTA5, HPGDS |
| PFDA | Stress response: detoxification: GST | *gst-9* | GSTA4, GSTA5, HPGDS |
| PFDA | Stress response: detoxification: GST | *gsto-2* | GSTO1, GSTO2 |
| PFDA | Stress response: detoxification: ugt | *ugt-1* | UGT3A1, UGT3A2 |
| PFDA | Stress response: detoxification: ugt | *ugt-16* | UGT3A1, UGT3A2 |
| PFDA | Stress response: detoxification: ugt | *ugt-19* | UGT3A1, UGT3A2 |
| PFDA | Stress response: detoxification: ugt | *ugt-2* | UGT3A1, UGT3A2 |
| PFDA | Stress response: detoxification: ugt | *ugt-20* | UGT3A1, UGT3A2 |
| PFDA | Stress response: detoxification: ugt | *ugt-24* | UGT3A1, UGT3A2 |
| PFDA | Stress response: detoxification: ugt | *ugt-41* | UGT3A1, UGT3A2 |
| PFDA | Stress response: detoxification: ugt | *ugt-61* | UGT1A6, UGT1A5, UGT3A2 |
| PFDA | Stress response: detoxification: ugt | *ugt-62* | UGT1A6, UGT1A5, UGT3A2 |
| PFDA | Stress response: detoxification: ugt | *ugt-7* | UGT3A1, UGT3A2 |
| PFDA | Transmembrane transport: ABC | *mrp-2* | ABCC1, ABCC2, ABCC3 |
| PFDA | Transmembrane transport: ABC | *pgp-1* | ABCB1, ABCB11, ABCB4 |
| PFDA | Transmembrane transport: ABC | *pgp-9* | ABCB1, ABCB11, ABCB4 |
| PFDA | Transmembrane transport: solute carrier | *C35A5.3* | SLC17A7, SLC17A1, SLC17A6, SLC17A8, SLC17A2, SLC17A4 |
| PFDA | Transmembrane transport: solute carrier | *C51E3.6* | SLC23A1, SLC23A2, SLC23A3 |
| PFDA | Transmembrane transport: solute carrier | *C53B4.3* | SLC22A18 |
| PFDA | Transmembrane transport: solute carrier | *T11G6.3* | SLC37A3 |
| PFNA | Stress response: detoxification: CYP | *cyp-13A6* | TBXAS1, CYP3A7, CYP3A5, CYP3A4 |
| PFNA | Stress response: detoxification: CYP | *cyp-13A7* | TBXAS1, CYP3A7, CYP3A5, CYP3A4 |
| PFNA | Stress response: detoxification: CYP | *cyp-34A10* | CYP21A2, CYP2U1, CYP2J2, CYP2R1 |
| PFNA | Stress response: detoxification: CYP | *cyp-34A9* | CYP21A2, CYP2U1, CYP2J2, CYP2R1 |
| PFNA | Stress response: detoxification: GST | *gst-33* | GSTA4, GSTA5, HPGDS |
| PFOS | Metabolism: short chain dehydrogenase | *C55A6.4* | CBR1 |
| PFOS | Metabolism: short chain dehydrogenase | *D2063.1* | SORD |
| PFOS | Metabolism: short chain dehydrogenase | *dhs-14* | HSD17B14 |
| PFOS | Stress response: detoxification: CYP | *cyp-13A12* | CYP3A7-CYP3A51P |
| PFOS | Stress response: detoxification: CYP | *cyp-13A4* | TBXAS1, CYP3A7, CYP3A5, CYP3A4 |
| PFOS | Stress response: detoxification: CYP | *cyp-13A5* | TBXAS1, CYP3A7, CYP3A5, CYP3A4 |
| PFOS | Stress response: detoxification: CYP | *cyp-13A6* | TBXAS1, CYP3A7, CYP3A5, CYP3A4 |
| PFOS | Stress response: detoxification: CYP | *cyp-13A7* | TBXAS1, CYP3A7, CYP3A5, CYP3A4 |
| PFOS | Stress response: detoxification: CYP | *cyp-34A10* | CYP21A2, CYP2U1, CYP2J2, CYP2R1 |
| PFOS | Stress response: detoxification: CYP | *cyp-34A9* | CYP21A2, CYP2U1, CYP2J2, CYP2R1 |
| PFOS | Stress response: detoxification: GST | *gst-5* | GSTA4, GSTA5, HPGDS |
| PFOS | Stress response: detoxification: GST | *gst-8* | GSTA4, GSTA5, HPGDS |
| PFOS | Stress response: detoxification: ugt | *ugt-1* | UGT3A1, UGT3A2 |
| PFOS | Stress response: detoxification: ugt | *ugt-16* | UGT3A1, UGT3A2 |
| PFOS | Stress response: detoxification: ugt | *ugt-19* | UGT3A1, UGT3A2 |
| PFOS | Stress response: detoxification: ugt | *ugt-20* | UGT3A1, UGT3A2 |
| PFOS | Stress response: detoxification: ugt | *ugt-41* | UGT3A1, UGT3A2 |
| PFOS | Stress response: detoxification: ugt | *ugt-61* | UGT1A6, UGT1A5, UGT3A2 |
| PFOS | Stress response: detoxification: ugt | *ugt-62* | UGT1A6, UGT1A5, UGT3A2 |
| PFOS | Stress response: detoxification: ugt | *ugt-7* | UGT3A1, UGT3A2 |
| PFOS | Transmembrane transport: ABC | *pgp-1* | ABCB1, ABCB11, ABCB4 |
| PFOS | Transmembrane transport: solute carrier | *C35A5.3* | SLC17A7, SLC17A1, SLC17A6, SLC17A8, SLC17A2, SLC17A4 |
| PFOS | Transmembrane transport: solute carrier | *C53B4.3* | SLC22A18 |
| PFOS | Transmembrane transport: solute carrier | *vglu-3* | SLC17A6, SLC17A7, SLC17A8 |

**Table S3.** Cross-species PFAS-responsive xenobiotic detoxification genes and corresponding literature evidence.

| Chemical Name | Cas RN | Gene Symbol | Organism | Interaction | PubMed IDs |
| --- | --- | --- | --- | --- | --- |
| PFDA | 335-76-2 | ABCB4 | *Mus musculus* | increases expression | 38181850 |
| PFDA | 335-76-2 | ABCC2 | *Mus musculus* | increases expression | 27344344\|38181850 |
| PFDA | 335-76-2 | ABCC2 | *Mus musculus* | affects reaction\|increases expression | 27344344 |
| PFDA | 335-76-2 | ABCC3 | *Mus musculus* | affects reaction\|increases expression | 18757529 |
| PFDA | 335-76-2 | ABCC3 | *Mus musculus* | increases expression | 18757529\|38181850 |
| PFDA | 335-76-2 | ABCC3 | *Mus musculus* | affects reaction\|increases expression | 18757529 |
| PFDA | 335-76-2 | CYP3A4 | *Homo sapiens* | increases expression | 23567314 |
| PFDA | 335-76-2 | UGT1A5 | *Mus musculus* | increases expression | 27344344 |
| PFDA | 335-76-2 | UGT1A5 | *Mus musculus* | affects reaction\|increases expression | 27344344 |
| PFNA | 375-95-1 | CYP3A4 | *Homo sapiens* | increases expression | 32588087 |
| PFNA | 375-95-1 | CYP3A4 | *Homo sapiens* | affects cotreatment\|increases expression | 38117326 |
| PFOS | 1763-23-1 | ABCB11 | *Homo sapiens* | increases expression | 21723365\|38199259 |
| PFOS | 1763-23-1 | ABCB11 | *Rattus norvegicus* | increases expression | 18692542 |
| PFOS | 1763-23-1 | ABCB4 | *Homo sapiens* | increases expression | 23567314 |
| PFOS | 1763-23-1 | CBR1 | *Mus musculus* | affects cotreatment\|increases expression | 36331819 |
| PFOS | 1763-23-1 | CBR1 | *Mus musculus* | affects cotreatment\|increases expression | 36331819 |
| PFOS | 1763-23-1 | CBR1 | *Mus musculus* | affects cotreatment\|increases expression | 36331819 |
| PFOS | 1763-23-1 | CBR1 | *Oncorhynchus mykiss* | increases expression | 21984479 |
| PFOS | 1763-23-1 | CYP2J2 | *Cyprinus carpio* | increases expression | 19162173 |
| PFOS | 1763-23-1 | CYP2J2 | *Cyprinus carpio* | increases expression | 18501439 |
| PFOS | 1763-23-1 | CYP2R1 | *Mus musculus* | affects cotreatment\|increases expression | 36331819 |
| PFOS | 1763-23-1 | CYP3A4 | *Homo sapiens* | affects cotreatment\|increases expression | 38117326 |
| PFOS | 1763-23-1 | CYP3A4 | *Homo sapiens* | affects cotreatment\|increases expression | 38117326 |
| PFOS | 1763-23-1 | CYP3A4 | *Homo sapiens* | increases expression | 21723365\|23567314\|27153767\|32253466\|32588087\|33772556 |
| PFOS | 1763-23-1 | GSTA4 | *Mus musculus* | affects cotreatment\|increases expression | 36331819 |
| PFOS | 1763-23-1 | GSTA4 | *Mus musculus* | affects cotreatment\|increases expression | 36331819 |
| PFOS | 1763-23-1 | GSTA4 | *Mus musculus* | affects cotreatment\|increases expression | 36331819 |
| PFOS | 1763-23-1 | GSTA4 | *Mus musculus* | increases expression | 26178269 |
| PFOS | 1763-23-1 | GSTA5 | *Homo sapiens* | increases expression | 33772556 |
| PFOS | 1763-23-1 | GSTA5 | *Mus musculus* | affects cotreatment\|increases expression | 36331819 |
| PFOS | 1763-23-1 | GSTA5 | *Mus musculus* | affects cotreatment\|increases expression | 36331819 |
| PFOS | 1763-23-1 | GSTA5 | *Rattus norvegicus* | increases expression | 19162173 |
| PFOS | 1763-23-1 | HSD17B14 | *Mus musculus* | increases expression | 26178269 |
| PFOS | 1763-23-1 | SLC17A8 | *Homo sapiens* | affects cotreatment\|increases expression | 38603627 |
| PFOS | 1763-23-1 | SLC22A18 | *Mus musculus* | affects cotreatment\|increases expression | 36331819 |
| PFOS | 1763-23-1 | SLC22A18 | *Mus musculus* | affects cotreatment\|increases expression | 36331819 |
| PFOS | 1763-23-1 | SLC22A18 | *Mus musculus* | affects cotreatment\|increases expression | 36331819 |
| PFOS | 1763-23-1 | SORD | *Mus musculus* | affects cotreatment\|increases expression | 36331819 |
| PFOS | 1763-23-1 | UGT1A5 | *Mus musculus* | affects cotreatment\|increases expression | 36331819 |
| PFOS | 1763-23-1 | UGT1A6 | *Rattus norvegicus* | increases expression | 19162173 |
